# An unprecedented DNA recognition–mimicry switch governs induction in arbitrium phages

**DOI:** 10.1101/2025.11.27.690993

**Authors:** Cora Chmielowska, Sara Zamora-Caballero, Javier Mancheño-Bonillo, Yuyi Li, Daniel Sin, Tom Borenstein, Shira Omer Bendori, Avigdor Eldar, Alberto Marina, José R Penadés

**Affiliations:** Department of Infectious Disease, Imperial College London, SW7 2AZ, London, UK; Centre for Bacterial Resistance Biology, Imperial College London, SW7 2AZ, London, UK; Instituto de Biomedicina de Valencia (IBV)-CSIC and CIBER de Enfermedades Raras (CIBERER)-ISCIII, 46010 Valencia, Spain; Shmunis School of Biomedicine and Cancer Research, Faculty of Life Sciences, Tel-Aviv University, Tel-Aviv, 69978, Israel; School of Health Sciences, Universidad CEU Cardenal Herrera, CEU Universities, Alfara del Patriarca, 46115, Spain

**Keywords:** repressor, activator, SOS response, Lysis-lysogeny decision, DNA damage, quorum sensing, temperate phages

## Abstract

Temperate phages integrate multiple information sources to regulate lysis-lysogeny transitions. SPBeta-like phages use arbitrium signalling and DNA damage to control repressor activity during lytic induction, but how the repressor functions and is inactivated by the SOS response remains unclear. Here, we show that SroF, the SPBeta-like phage repressor, binds DNA via a novel mechanism involving its integrase-like fold, enabling stable prophage repression. Upon DNA damage, the host SOS response triggers derepression of a newly identified antirepressor, Sar. Sar binds SroF by mimicking the DNA structure recognised by the repressor, inactivating its function and inducing phage. This mechanism is conserved across SPβ-like phages, which encode multiple, specific SroF-Sar pairs. Surprisingly, repressor inactivation alone is insufficient for induction when arbitrium levels are high. Our results uncover the mechanism underlying the double layer of control that ensures phage induction occurs only under SOS conditions and in the absence of neighbouring prophages.

## Introduction

Bacteriophages (phages) are viruses that infect bacteria. Upon infection, temperate phages can either multiply and kill the bacterial host through the lytic cycle or enter the lysogenic (prophage) state, in which they typically integrate into the bacterial chromosome and replicate passively alongside the host. Dormant prophages may later reactivate (induce) under certain conditions, re-entering the lytic cycle.^1–6^

*Escherichia coli* phage Lambda is the classic model organism for studying how temperate phages alternate between lytic and lysogenic life cycles.^7^ To establish and maintain lysogeny, phage Lambda relies on the CI master repressor protein, which inhibits the expression of lytic genes. The CI lambda repressor comprises two functional domains: an N-terminal domain that binds specific DNA sequences to block transcription, and a C-terminal domain that mediates dimerization and cooperative interactions essential for stable repression and autoregulation.^8^

Since prophage survival depends on the survival of their bacterial hosts, induction of many prophages, including phage Lambda, is linked to the bacterial SOS response, a global regulatory system that is activated in response to DNA damage.^9,10^ It serves as a coordinated defence mechanism to repair DNA lesions and ensure bacterial survival. During normal growth, the LexA protein represses the expression of bacterial genes comprising the SOS regulon by binding to a DNA regulatory motif known as the SOS box.^10,11^ Upon DNA damage, the bacterial RecA protein is activated and stimulates LexA’s self-cleavage.^12,13^ In phage Lambda, the CI repressor contains a LexA-like domain and also undergoes RecA-mediated autoproteolysis during the SOS response.^9,13,14^ Cleavage of CI abolishes oligomerisation and, consequently, repression, allowing expression of lytic genes and switching the phage from lysogeny to the lytic cycle.

While CI-like repressors are widespread among temperate phages infecting diverse bacterial species, alternative phage induction mechanisms have been discovered. These include direct LexA regulation of lytic genes or inactivation of phage repressors by small phage-encoded antirepressor proteins.^15–20^ The expression of these antirepressors can be regulated by factors such as LexA or host quorum-sensing signals.^16–20^

Another distinct mechanism of lysis-lysogeny control has been reported in a family of *Bacillus* phages that utilise the arbitrium system. Unlike other systems, these temperate phages base their decision between the lysogenic and lytic cycles not only on the physiological state of the host bacterium but also on the anticipated availability of future susceptible hosts. To guide this decision, arbitrium phages encode a quorum-sensing-like communication system that operates during infection and induction.^21–24^ This small-peptide-based signalling mechanism enables *Bacillus* phages to coordinate their life-cycle choices collectively by sensing phage population density as a proxy for host availability. During infection, arbitrium phages produce a peptide called AimP. Early in infection, when bacterial cells are abundant relative to phages, AimP is present at low levels, favouring the lytic cycle. However, after multiple infection rounds, as phage numbers rise and bacterial concentrations decline, elevated AimP levels promote lysogeny. At this stage, arbitrium phages express a six-gene operon named the “<u>S</u>PBeta repressor <u>o</u>peron” (*sroA*–*sroF*), which includes the master repressor SroF.^23,25,26^

Importantly, the arbitrium repressor SroF is structurally and functionally distinct from any previously characterised phage repressor. In the prototypical SPBeta phage, SroF displays a unique integrase/tyrosine recombinase fold not found in other known phage repressors.^25,27^ Unlike Lambda CI, which functions through cooperative binding of multiple CI dimers to palindromic operator sequences, SroF recognises a considerably longer DNA motif with multiple binding sites distributed across distant regions of the phage genome.^7,25,27^ How this binding occurs, however, remains to be determined. Moreover, the mechanism by which SroF is inactivated during the SOS response differs fundamentally from Lambda CI. Although arbitrium prophages are induced by the SOS response, SroF lacks a LexA-like domain. These significant differences between the well-studied CI-like repressors and SroF present a compelling model for exploring alternative modes of phage repressor function and regulation.

This study aimed to elucidate the molecular mechanism of the unique lysis-lysogeny switch encoded by the SPBeta family of arbitrium phages and to provide new insights into its regulation. We focused on understanding how SroF repressors bind DNA and how their activity is inactivated in response to the SOS response. Contrary to most other repressors, we found that SroF binds DNA as monomers, recognising up to 24 nucleotides of its DNA motif. Release of SroF from DNA does not involve proteolytic cleavage but instead occurs through interaction with a small antirepressor protein, which we named Sar (<u>S</u>PBeta-family <u>a</u>ntirepressor). Bioinformatic analyses indicate that this repressor–antirepressor system is conserved across other SPBeta-family phages, a finding we experimentally confirmed for two arbitrium model phages, Phi3T and SPBeta. Completing the picture, we demonstrate that antirepressor expression is directly controlled by the SOS response. Furthermore, we show that the SOS signal is integrated upstream of arbitrium-mediated induction control, revealing that lysogen survival under stress balances information from both host stress and prophage population density. Altogether, our results reveal a sophisticated and unprecedented regulatory network that finely tunes the lysis-lysogeny decision, representing a major advance in our understanding of phage biology.

## Results

### Structural characterization of SroF bound to DNA

In previous publications we characterised the apo form of the SroF master repressor for phage SPBeta (SroF^SPBeta^), showing it adopts a monomeric integrase/recombinase-like fold, unique among phage repressors.^25^ To further investigate the molecular basis of this novel mechanism of repression in the SPBeta phage family, we solved the structure of the SroF repressor bound to its DNA box. In this study, we used the SroF repressor from the model phage Phi3T (SroF^Phi3T^), which was originally used to describe the arbitrium system and its 26 bp DNA binding box (CAGTAATTTTATTTTTATTCTTAATG). This allowed us to confirm the conserved folding of the SroF proteins, and to investigate their mechanism of DNA binding. The crystals of SroF^Phi3T^-DNA complex diffracted up to 2.5 Å resolution (Supplementary Table S1) and showed in its asymmetric unit the presence of one SroF repressor and one DNA box. Unlike other integrases/recombinases that recognise DNA as a dimer and form tetramers during the recombination process, the presence of biological oligomers produced by crystallographic symmetry was discarded, confirming that SroF acts as a monomeric repressor. In the structure of the complex, SroF^Phi3T^ displays the integrase-like fold with an all-helical N-terminal Core Binding (CB) domain (residues 1-105) and a C-terminal Catalytic (CAT) domain (residues 113-319) that are joined by an unstructured connector (Supplementary Figure S1A). SroF^Phi3T^ (solved in a complex with DNA) and SroF^SPBeta^ (solved in apo form) showed very similar folding (RMSDs of 1.8 Å and 2.4 Å for the comparison of the CB and CAT domains respectively), indicating that the repression mechanism would be conserved in both phages, but strong conformational changes are observed between the apo and DNA bound structure. To bind the DNA, the N-terminal CB domain of the repressor approaches the C-terminal CAT domain, a movement facilitated by the flexible interdomain linker, adopting a closed conformation that nearly encircles the DNA (Figure 1A). This movement, which constitutes the more visible effect on the protein upon binding to DNA, creates a C-shaped cavity, with a positively charged inner surface, while the outer surface predominantly exhibits neutral or negative charges (Figure 1B).

**Figure 1:**
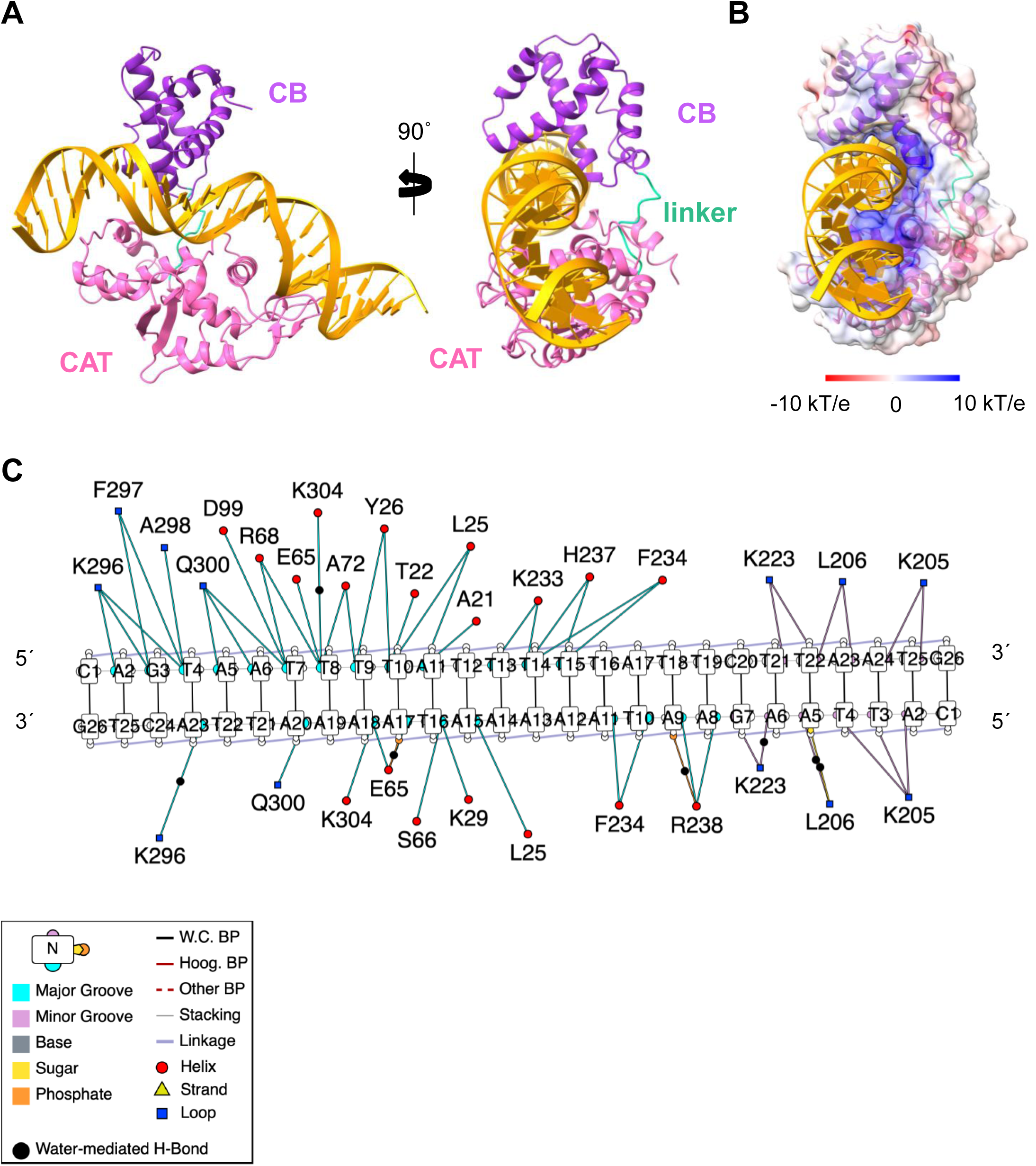
Crystal structure of SroF^Phi3T^ in complex with DNA. (**A**) Two views of a ribbon representation of the crystal structure of SroF^Phi3T^-DNA complex. DNA is shown in orange. SroF^Phi3T^ core domain (CB) is coloured in purple. Catalytic domain (CAT) is depicted in pink whereas the unstructured linker is shown in turquoise. (**B**) SroF^Phi3T^ electrostatic surface potential as calculated by APBS and visualized in ChimeraX. Key indicates the charge of the surface. (**C**) DNA readout by SroF^Phi3T^ as represented by DNAproDB. Contacting residues are indicated for each DNA base. Legend indicates the colour code for the interactions and the secondary structure element where the contacting amino acids are located.

Unlike other integrases/recombinases, which tend to make a limited number of base-specific contacts^28^, SroF performs extensive direct read-out of its DNA operator, interacting with 24 out of 26 bp in the box (only the 3’ and 5’ bases at the ends are not read) (Figure 1C and Supplementary Figure S2A). The CAT domain accommodates most of the DNA, recognising the box through interactions in both the major and minor grooves. On one side, the CAT domain introduces helix α_13_ across one DNA major groove, recognising the DNA sequence using K233, F234, H237 and R238 residues. In the same vein, helix α_18_ is placed in the major groove in the following DNA turn of the operator, establishing specific contacts with the DNA bases through Q300 and K304. On the opposite side of the operator, DNA recognition occurs instead through the minor groove, involving residues L205, L206 or K233, among others, located in the loops α_12_-β_4_ and β_4_-α_13_. Meanwhile, the CB domain bends over a 6 bp central part of the box, inserting its helix α2 in a DNA major groove and using the amino acids T22, L25 and Y26 to read out the sequence (Figure 1C, Supplementary Figure S1A and Supplementary Table S2). Besides the aforementioned specific interactions, multiple contacts are established with the DNA backbone by both the CB and CAT domains (Supplementary Figure S2A and Supplementary Table S2). As a result, SroF^Phi3T^ is able to establish contacts along the entire DNA molecule (Figure 1C and Supplementary Figure S2A).

The DNA section recognised through the major groove by the two SroF domains (bases 11 to 20) corresponds to the two consecutive A-tracts (AAAAATAAAA), a type of sequence that has been described to have narrower minor grooves and to bend the DNA.^29–32^ Analysis of DNA topology in the structure with DNAproDB^33^ confirms the narrowing of the minor groove up to a width of ∼ 2.8 Å. Therefore, it is tempting to speculate that besides the DNA sequence, SroF^Phi3T^ is also recognizing the shape of the DNA. Such dependence on DNA shape seems to be related to the folding of SroF, since similar recognition of DNA conformation has been found in other integrases/recombinases such as the integron integrase or Cre recombinases.^34–36^

The SroF^Phi3T^-DNA structure shows a striking lack of resemblance to the lambda CI-DNA structure. Unlike the dimeric lambda repressor, which binds to the DNA only by the insertion of a single alpha helix with an interface area of ∼420 Å², SroF^Phi3T^ interacts as a monomer, encircling the DNA, interacting with multiple bases and increasing its interface area to ∼1700 Å². The CI repressor of phage lambda is a functional homologue of LexA, which undergoes RecA-mediated autocleavage upon activation of the bacterial host SOS response, thereby leading to prophage induction.^9,13,14^ However, SroF^Phi3T^ lacks the characteristics necessary for proteolytic cleavage, and, if cleaved, the resulting proteolysis would be insufficient to release SroF^Phi3T^ from DNA given the extensive binding interfaces on both SroF^Phi3T^ domains. Therefore, we hypothesised that a different mechanism would be required to facilitate the DNA release of this integrase-like repressor.

### The phage-encoded phi3T_99 protein directly interacts with SroF^Phi3T^

To gain further insight into how this atypical, integrase-like repressor system functions, we conducted protein pull-down assays to find direct interaction partners of SroF^Phi3T^. We hypothesised that, for the prophage to be activated, a phage- or bacterially-encoded protein whose expression is controlled by the SOS response would bind to the SroF repressor to inactivate its function. The pull-down assay was conducted using a *Bacillus subtilis* 168 Δ6 strain lysogenised with Phi3T and an ectopically expressed a 3xFLAG-tagged version of SroF^Phi3T^. Two independent pull-down assays were analysed by mass spectrometry, yielding several phage candidate proteins (Supplementary Table S3). However, only the phi3T_99 protein, of unknown function, appeared in both replicates. Interestingly, the gene encoding this small protein is located one open reading frame (ORF) downstream of the master repressor gene *sroF*^phi3T^ (Figure 2A). Using biolayer interferometry (BLI), we validate the direct interaction between recombinant SroF^Phi3T^ and Phi3T_99, showing high affinity binding in the low nanomolar range (K_D_ 21.7 nM) (Supplementary Figure S2B).

**Figure 2.**
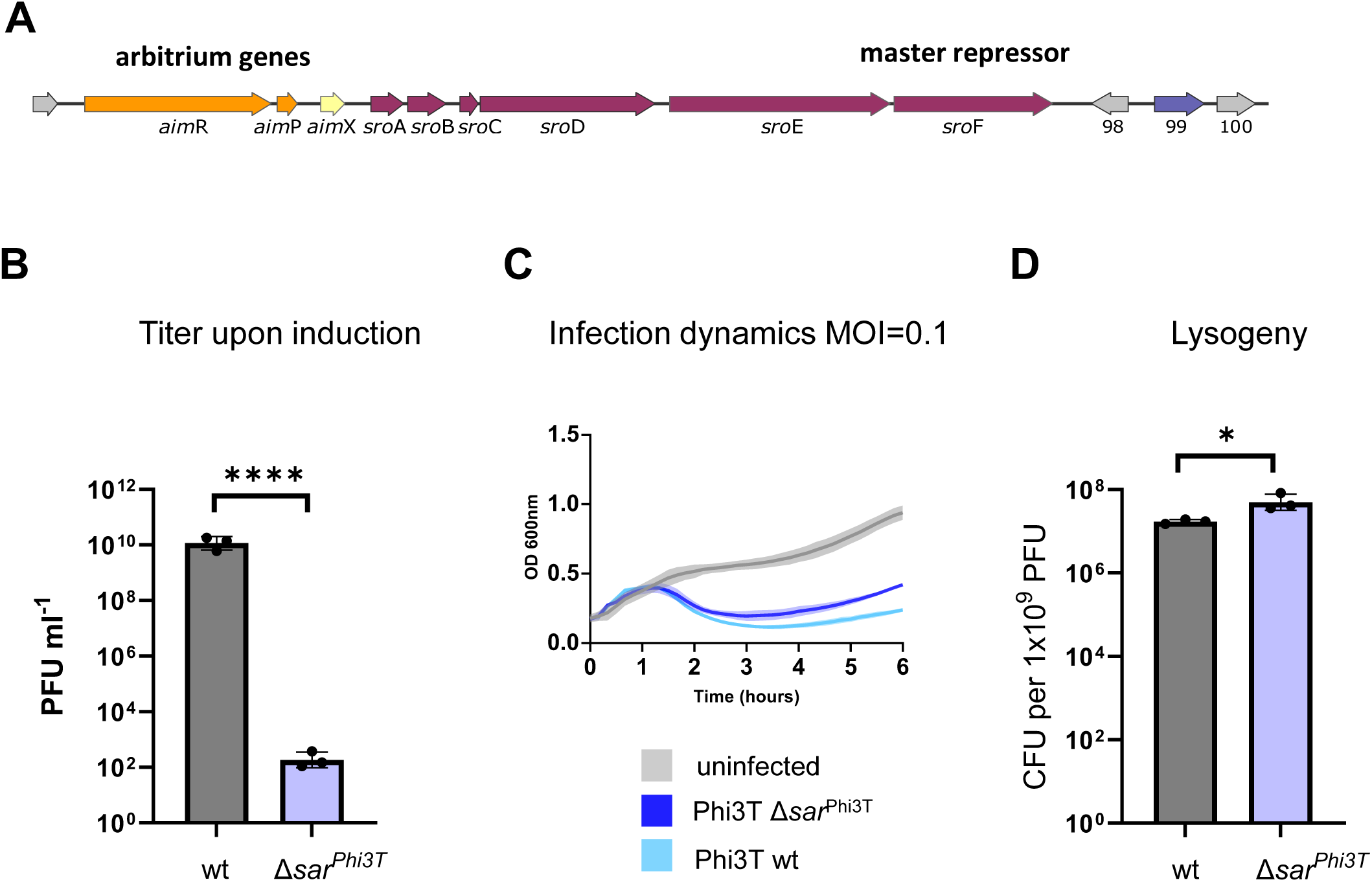
Sar^Phi3T^ (Phi3T_99) is necessary for efficient phage induction. **(A)** Schematic overview of the arbitrium operon (orange), 6-gene operon (pink) and *phi3T_99*/*sar*^Phi3T^ (purple) region of phage Phi3T. (**B**) Strains lysogenic with phages Phi3T wt or Phi3T Δ*sar*^Phi3T^ were MMC induced. Resulting phage particles were quantified using 168 Δ6 as a recipient strain. (**C**) Strain 168 Δ6 was infected with phages Phi3T wt or Phi3T Δ*sar*^Phi3T^ at MOI=0.1 and infection dynamics were followed by measuring OD_600_ readings. (**D**) The number of lysogens resulting from an infection of 168 Δ6 by Phi3T wt or Phi3T Δ*sar*^Phi3T^ phages. The results are shown in panel (**B**) as PFUs ml^-1^ and in panel (**D**) as CFUs ml^−1^ normalised by PFUs ml^-1^ and represented as the CFU of an average phage titer (1×10^9^ PFU ml^-1^). Data represents geometric mean and SD (n=3) in panels (**B**) and (**D**) and the mean and SD (n=3) in panel (**C**). Two-tailed t-tests were performed. p-values are indicated above each comparison: ∗∗∗∗ p < 0.0001; ∗ p < 0.05.

### Phi3T_99 (Sar^Phi3T^) is the antirepressor protein

To test the hypothesis that Phi3T_99 (hereafter referred to as Sar^Phi3T^) could be an antirepressor protein whose expression inactivates SroF function, we generated a deletion mutant of this gene in the Phi3T prophage, which also carries a kanamycin resistance marker. The wild-type (wt) and *sar*^Phi3T^mutant prophages were then induced with Mitomycin C (MMC), which activates the SOS response, and phage titres were quantified. Notably, deletion of the ORF encoding Sar^Phi3T^ severely impaired induction of the mutant phage compared to the wt, resulting in a 10^8^-fold reduction in the number of infectious particles (Figure 2B). These results suggest that Sar^Phi3T^ is required for efficient SOS-mediated induction.

Next, we studied the effect of *sar*^Phi3T^ deletion on infection by measuring the growth curve of naïve Δ6 *B. subtilis* liquid cultures infected with wt *or* Δ*sar*^Phi3T^ Phi3T phages. We observed a slightly higher bacterial survival in infections with the deletion mutants than wt phages (Figure 2C). We also conducted lysogeny assays and found that the number of lysogens (CFU per PFU) produced when infecting with Phi3T Δ*sar*^Phi3T^ phages was only slightly higher compared to those generated by wt Phi3T (Figure 2D). Therefore, the Δ*sar*^Phi3T^ phage seemed to be defective in the SOS induction process to a much higher degree than the infection process. All these results are in accordance with the idea that Sar^Phi3T^ is the arbitrium antirepressor protein, which facilitates the release of the repressor SroF from DNA. Therefore, we renamed Phi3T_99 as Sar^Phi3T^ (from **S**Pbeta-family **a**nti**r**epressor).

To confirm whether Sar^Phi3T^ is indeed an antirepressor protein, we hypothesised that its expression would be sufficient to activate the lytic cycle of the prophage in the absence of an SOS response. To test this, we cloned the *sar*^Phi3T^ gene into an integrative expression vector pDR110 under the control of an Isopropyl ß-D-1-thiogalactopyranoside (IPTG) inducible promoter. Next, we transformed the integration vector into a strain harbouring the Phi3T Δ*sar*^Phi3T^ prophage. When Sar^Phi3T^ was expressed upon adding IPTG, the bacteria lysed, as observed by following growth dynamics in liquid cultures (Figure 3A). This was observed independent of whether MMC was present or absent. The expression of Sar^Phi3T^ in strains without prophages did not lead to any growth defects, showing that the observed lysis is phage dependent (Supplementary Figure S3). In contrast, the strain harbouring a wt prophage and empty control vector required MMC for lysis (Figure 3B), while Phi3T Δ*sar*^Phi3T^prophages carrying the empty control vectors lysed poorly independent of the presence of MMC (Figure 3C). We next confirmed that the observed cell lyses was due to phage induction by counting the number of resulting phage particles in plaque assays (Figure 3D). Therefore, expression of Sar^Phi3T^ from an expression vector is sufficient to cause prophage induction, independently of the SOS response.

**Figure 3.**
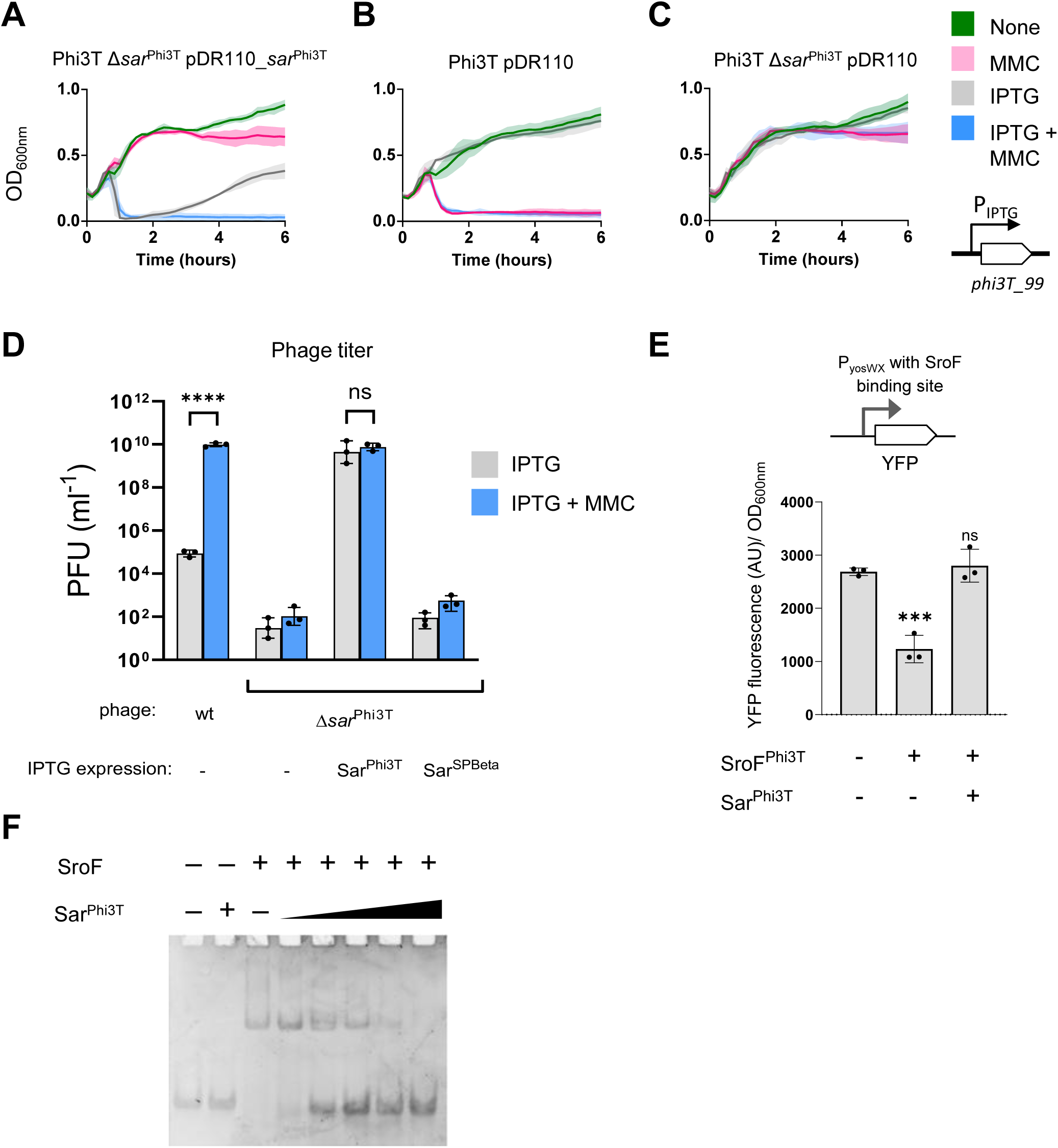
Expression of Sar^Phi3T^ in lysogens leads to phage induction in a MMC-independent way by interfering with SroF function. (**A-D**) Strains lysogenic with phages Phi3T wt or Phi3T Δ*sar*^Phi3T^ were complemented with either empty integration vector (amyE::Pspank-) or with IPTG-inducible *sar*^Phi3T^ or *sar*^SPBeta^. IPTG (gray), MMC (pink) or both IPTG and MMC (blue) were added to the cultures and cell lysis was followed by measuring OD_600_ readings. Means and SD’s are presented (n = 3). (**D**) The number of resulting phages upon addition of IPTG (gray) or IPTG and MMC (blue) was quantified using 168 Δ6 as a recipient strain. The results are shown as PFUs ml^-1^, with geometric means and SDs presented (n = 3). A two-tailed t-test on log_10_ transformed data was performed to compare mean differences. (**E**) Sar^Phi3T^ interferes with SroF repression of YFP transcriptional reporter constructs. Expression of YFP from yosXY promoter (containing a Phi3TRE box recognized by SroF) was measured in the presence or absence (as indicated by the +/- symbols under the x-axis) of xylose-inducible *sro*F and/or IPTG-inducible *sar*^Phi3T^. Arbitrary fluorescence unit measurements normalized by OD_600_ readings are shown, with means and SDs presented (n = 3). An ordinary one-way ANOVA of followed by a Dunnett’s multiple comparisons test was performed to compare mean differences with the control strain not expressing SroF or Sar^Phi3T^. (**F**) Electromobility shift assay to assess the competition for SroF binding site between the dsDNA probe and Sar^Phi3T^ (gradient of concentrations). p values are indicated above the plots: p ≥ 0.05 not significant (ns), p < 0.001 (∗∗∗).

Next, we verified if the observed phage induction was a result of a Sar^Phi3T^ interference with SroF^Phi3T^ repression function. A reporter system was used, consisting of a yellow fluorescent protein (YFP) gene cloned under the control of one of the promoters regulated by SroF^Phi3T^ (Figure 3E). The reporter contains a SroF^Phi3T^ repressor binding box, Phi3TRE, as previously described.^25^ When SroF^Phi3T^ is present (induced from a Xylose promoter), YFP expression is reduced due to binding of SroF^Phi3T^ to the master repressor binding box. However, when we simultaneously expressed Sar^Phi3T^ (induced from a IPTG promoter) and SroF^Phi3T^, the high YFP expression was restored (Figure 3E).

Finally, we performed an *in vitro* competition assay of SroF^Phi3T^ between Sar^Phi3T^ and its DNA binding box and we evaluated the results via an electrophoretic mobility shift assay (EMSA). Upon incubation of the SroF^Phi3T^-DNA complex with increasing concentrations of Sar^Phi3T^, the DNA was displaced from SroF^Phi3T^, indicating that Sar^Phi3T^ disrupts the repressor:DNA complex, releasing DNA from SroF^Phi3T^. At a SroF^Phi3T^:Sar^Phi3T^ molar ratio of 1:1 almost all the DNA was dissociated from SroF^Phi3T^ binding, and a total disappearance of SroF^Phi3T^-DNA complex was achieved with a SroF^Phi3T^:Sar^Phi3T^ ratio of 1:2 (Figure 3F). These results collectively indicate that Sar^Phi3T^ facilitates the release of SroF^Phi3T^ from the DNA, thereby inactivating the repressor’s function.

### Structural characterization of SroF-Sar complex

To gain deeper insight into the molecular basis on how Sar^Phi3T^ disrupts the interaction of SroF^Phi3T^ with its cognate DNA operator, we characterised the crystal structure of the Sar^Phi3T^- SroF^Phi3T^ complex. The crystal structure, solved at 3 Å resolution (Supplementary Table S1), showed in the asymmetric unit a complex comprised of a Sar^Phi3T^ homodimer that binds two SroF^Phi3T^ monomers (Figure 4A). In the homodimer, Sar^Phi3T^ is composed of four β-strands (β_1_-β_4_), three α-helices (α_1_ and α_3_) and an unstructured C-terminal portion of 7 amino acids (Supplementary Figure S1B). Interestingly, the Sar^Phi3T^ homodimer is an intertwined dimer nucleated for the antiparallel disposition of the long β_1_ strands (residues 4-21) surrounded by the β_4_ strands to form a core molecular β-sheet. While one side of this core β-sheet is exposed, the opposite side is decorated with the three α-helices and the β-hairpin (formed by β_2_-β_3_ strands) from each monomer (Figure 4A). The molecular intertwining gives rise to a highly extensive dimerization interface, with a buried surface area greater than 3500 Å², indicative of a stable and specific dimeric that offer symmetrical binding motifs in its flanks that recruit two independent SroF^Phi3T^ monomers.

**Figure 4:**
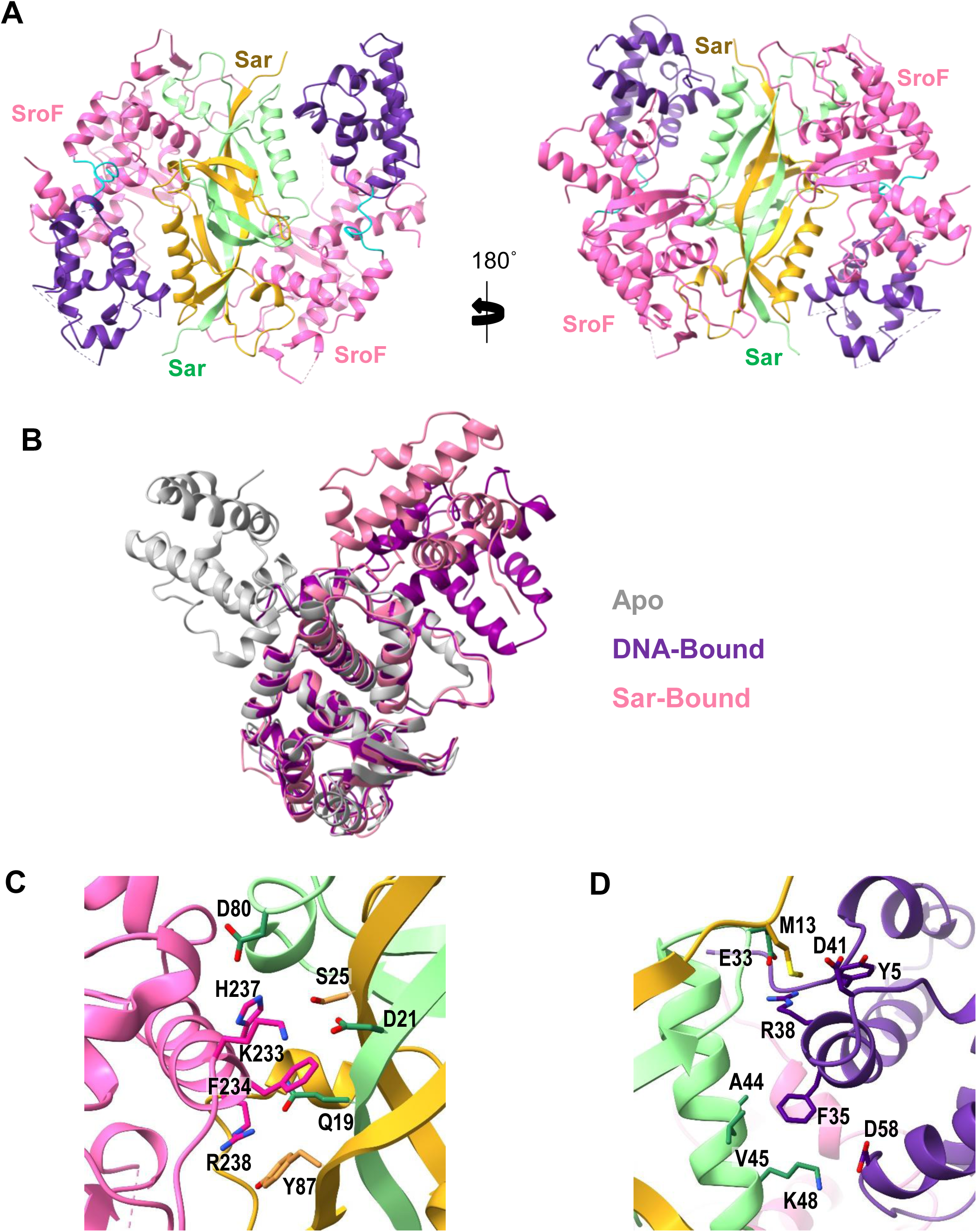
Crystal structure of SroF^Phi3T^-Sar^Phi3T^ complex shows a competition for SroF^Phi3T^ binding sites between Sar^Phi3T^ and DNA. (**A**) Two different views of a ribbon representation of the crystal structure of SroF^Phi3T^-Sar^Phi3T^ complex. SroF^Phi3T^ is depicted purple and magenta for the CB and CAT domains, respectively, and monomers of the Sar^Phi3T^ homodimer in yellow and pale green. (**B**) Ribbon representation of SroF^Phi3T^ CAT domain superposition of different SroF structures. In grey, apo SroF^SPBeta^, in pink SroF^Phi3T^ structure when bound to Sar^Phi3T^ and in purple SroF^Phi3T^ structure when bound to DNA. (**C**) Zoomed view of the contact area between SroF^Phi3T^ α13 helix CAT domain and (**D**) α1 helix CB domain. Interacting residues are shown in sticks and labelled. Molecules are coloured as in A

A superimposition of the CAT domains from SroF either in the absence of ligands or when bound to DNA or the antirepressor showed that Sar^Phi3T^ stabilises SroF in an intermediate conformation, positioned between the open apo and the closed DNA-bound states (Figure 4B). This superimposition also illustrates how the dimeric Sar^Phi3T^ interacts with SroF^Phi3T^, enabling it to outcompete DNA in SroF binding. Interestingly, Sar^Phi3T^ uses different interaction strategies with the two SroF domains. It mimics the DNA in order to bind to the CAT domain while shifting the CB domain to generate a new interaction surface with it. Sar^Phi3T^ mimics both the DNA sequence recognised by SroF^Phi3T^ interacting, among others, with helix α_13_ (i.e. K233, F234, H237 or R238) through its β_1_ (residues Q19, D21, S25, D80 or Y87) (Figure 4C), as well as emulating the DNA backbone via the mimicking of phosphate by multiple negatively charged residues (E79, D80 of one monomer and E89, E93, D96, E100, E101 on the other monomer) and additional interactions (Figure 4C, Supplementary Table S2). On the other hand, the interaction surface with the SroF^Phi3T^ CB domain is completely different from the one used to recognise DNA. It mainly involves residues at the N-terminal end (Y5), α_2_ (F35, R38, D41) and α_4_ (D58) of the repressor, which interact with E33, and α_1_ residues A44, V45, and K48 of one monomer, and with M13 of the second monomer of Sar^Phi3T^ dimer (Figure 4D). Although Sar^Phi3T^ does not interact with the residues involved in DNA recognition by the CB domain, the displacement and rotation of this domain induced by Sar occludes partially this surface, adding an additional impediment to SroF DNA recognition. Altogether the Sar^Phi3T^ dimer establishes an extensive interaction interface, exceeding 1800 Å² with each of the two repressors it binds. This remarkably large surface area is critical for enabling Sar to effectively compete with a DNA operator of substantial length (24 pb) and to cause efficient derepression, highlighting the structural and functional sophistication of the Sar-SroF interaction.

### Sar^Phi3T^ expression is controlled by LexA

The previous results demonstrated that the expression of Sar^Phi3T^ is responsible for the SOS-mediated Phi3T phage induction, leading to the question: how does the SOS response regulate Sar^Phi3T^ expression? Using the *B. subtilis* LexA-binding motif (SOS box) consensus sequence^37^, a bioinformatics search revealed the presence of a putative SOS box upstream of *sar*^Phi3T^ (Figure 5A), suggesting that the gene’s expression is directly regulated by LexA, the master SOS repressor.

**Figure 5.**
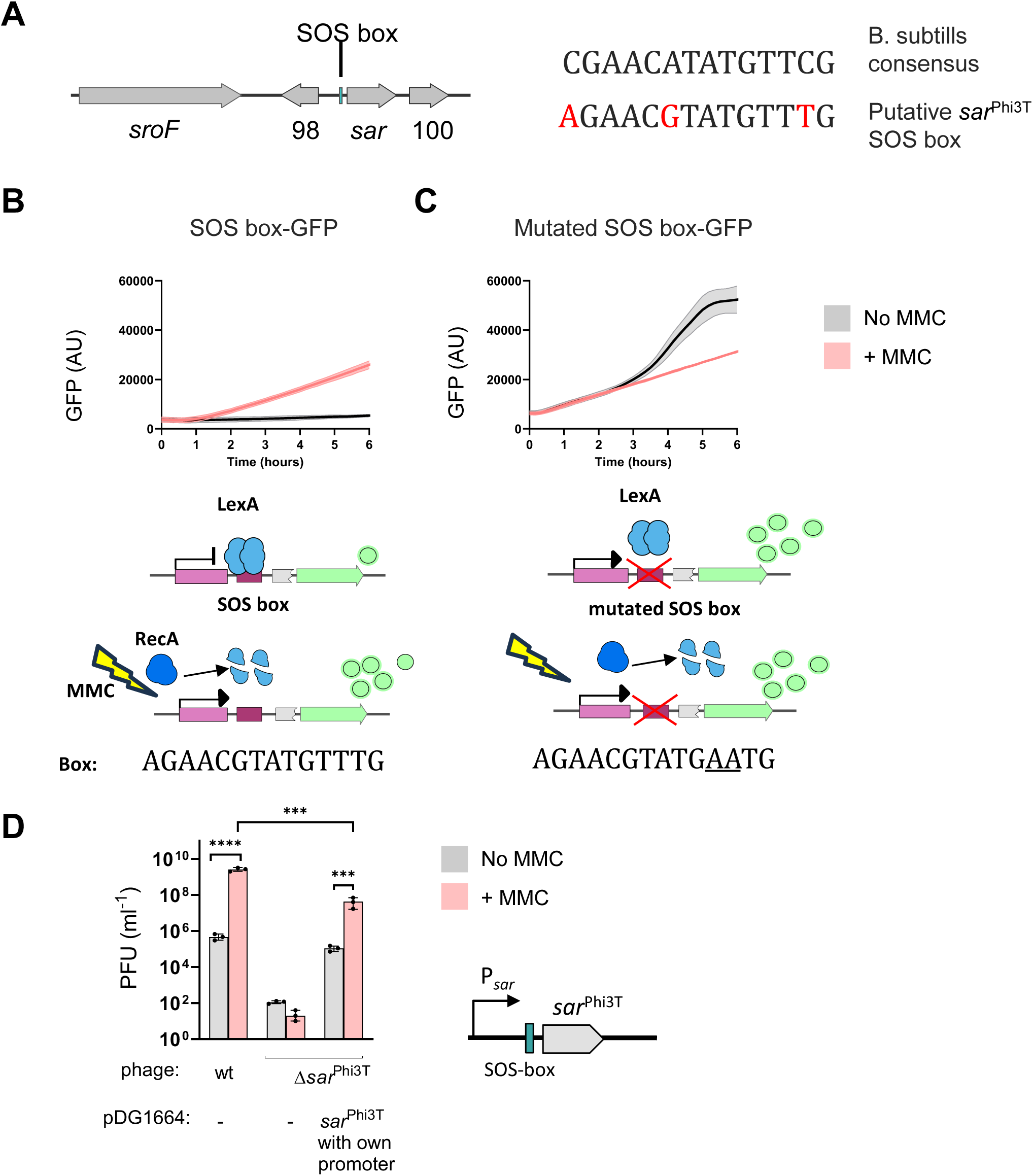
Sar^Phi3T^ expression is regulated by SOS response via LexA-binding. (**A**) Schematic overview of SOS box location upstream of *sar*^Phi3T^ gene. Comparison of consensus sequence for *B. subtilis* SOS-boxes and the putative SOS box upstream of *sar*^Phi3T^, with nucleotide differences marked in red. (**B-C**) Expression of GFP from transcriptional fusion with *sar*^Phi3T^ promoter was measured (arbitrary fluorescence units), with means and SDs presented (n = 3). The expression was measured either in absence (grey) or presence of MMC (pink). The constructs contained either an intact SOS box (**B**) or a version with a mutation in the conserved positions (underlined) (**C**). A schematic of constructs is presented: GFP (green), LexA (blue), RecA (dark blue) *sar*^Phi3T^ predicted promoter (pink), SOS box (dark pink). Due to the reduced growth of cells when treated with MMC, the fluorescence values were not normalised by OD measurements; however within a treatment group the strains had similar growth rates (Supplementary Figure S4). (**D**) Strains lysogenic with phages Phi3T wt or Phi3T Δ*sar*^Phi3T^ were complemented with either empty integration vector (pDG1664) or with construct containing *sar*^Phi3T^ gene with its own promoter and SOS box. The number of phages produced in presence or absence of MMC was quantified using 168 Δ6 as a recipient strain. The results are shown as PFUs ml^-1^, with geometric means and SDs presented (n = 3). A two-tailed t-test on log_10_ transformed data was performed to compare mean differences. p values are indicated above the plots: p < 0.001 (∗∗∗).p < 0.0001 (∗∗∗∗).

To test this, the *sar^Phi3T^* promoter was transcriptionally fused to a GFP (green fluorescent protein), cloned into an integrative vector and the expression of GFP was measured in the absence or presence of MMC (Figure 5B, Supplementary Figure S4). High GFP signals were only detected when Mitomycin C was added (Figure 5B). Next, a 2 nt point mutation was introduced into the predicted SOS box (Figure 5C). This mutation was previously shown to partially prevent LexA binding.^22^ When using the mutated SOS box construct, high GFP levels were produced independent of MMC presence, confirming that the SOS box is necessary for controlling the expression of Sar^Phi3T^ (Figure 5C). These results confirm that the expression of s*ar^Phi3T^* is regulated by the SOS response by binding of LexA to the upstream SOS box, thereby establishing a direct molecular link between the host SOS response and phi3T phage induction.

Next, we tested the functionality of this LexA control, and cloned *sar*^Phi3T^ together with the upstream region predicted to contain the natural promoter of Sar^Phi3T^ (together with the putative SOS box) into an integrative vector pDG1664 lacking an external expression promoter. The vector was then integrated into a strain harbouring a Phi3T Δ*sar^Phi3T^* prophage. Complementation with this vector restored SOS-dependant induction of the phage (Figure 5D). Notably, the complementation was only partial, and not to a wt phage level. This could possibly be explained by the presence of only 1 gene copy in the chromosome in complemented strains, in contrast to predicted multiple copies of *sar^Phi3T^* present in induced, replicating wt phages.

### Arbitrium system acts downstream Sar^Phi3T^ expression and increases survival of lysogens under stress conditions

To characterise the interaction between Sar^Phi3T^ and SroF, we hypothesised that overexpression of Sar^Phi3T^ in a Phi3T lysogen would impose a strong selective pressure for mutations that disrupt the Sar-SroF interaction. Such mutations would render the lysogen insensitive to the SOS response, allowing cells to survive phage induction. Specifically, we expected to recover cells carrying mutations either in *sar*^Phi3T^ or *sroF*^Phi3T^ that prevent activation of the lytic cycle.

To test this hypothesis, we used a Phi3T Δ*sar*^Phi3T^ lysogen in which two ectopic copies of *sar*^Phi3T^ were integrated into the chromosome under the control of xylose- and IPTG-inducible promoters. Following induction with both xylose and IPTG, cultures were plated to isolate rare survivors - cells in which the phage had not induced efficiently. These surviving colonies were sequenced to identify mutations that conferred resistance to phage induction. Surprisingly, none of the recovered mutants carried changes in *sroF*. Instead, all isolates harboured either frameshift mutations in the *aimR* gene, which encodes the receptor of the arbitrium system, or mutations in the AimR binding site (Supplementary Table S4). Both types of mutations likely impair the functionality of the arbitrium system.

AimR is an antiterminator whose activity is modulated by the quorum-sensing peptide AimP.^26^ In the absence of AimP, AimR activates expression of AimX, which promotes lytic development by interacting with different phage and cellular partners, including the bacterial toxin MazF.^26,38^ By contrast, when AimP is present, it binds to AimR and inhibits its activity, thereby repressing *aimX* expression and favouring lysogeny. In the absence of AimX, MazF degrades most phage mRNAs except those of a key six-gene operon that includes *sroF*, promoting lysogeny.^38^

These evolutionary experiments suggested that the arbitrium system functions downstream of *sar*^Phi3T^ expression to regulate phage induction. To test this model directly, we used a strain carrying a Phi3T Δ*sar*^Phi3T^ lysogen and an integrated IPTG-inducible *sar*^Phi3T^ gene. As expected, phage induction driven by IPTG-induced expression of Sar^Phi3T^ was strongly suppressed by the addition of synthetic arbitrium peptide (5 μM) (Figure 6A).

**Figure 6.**
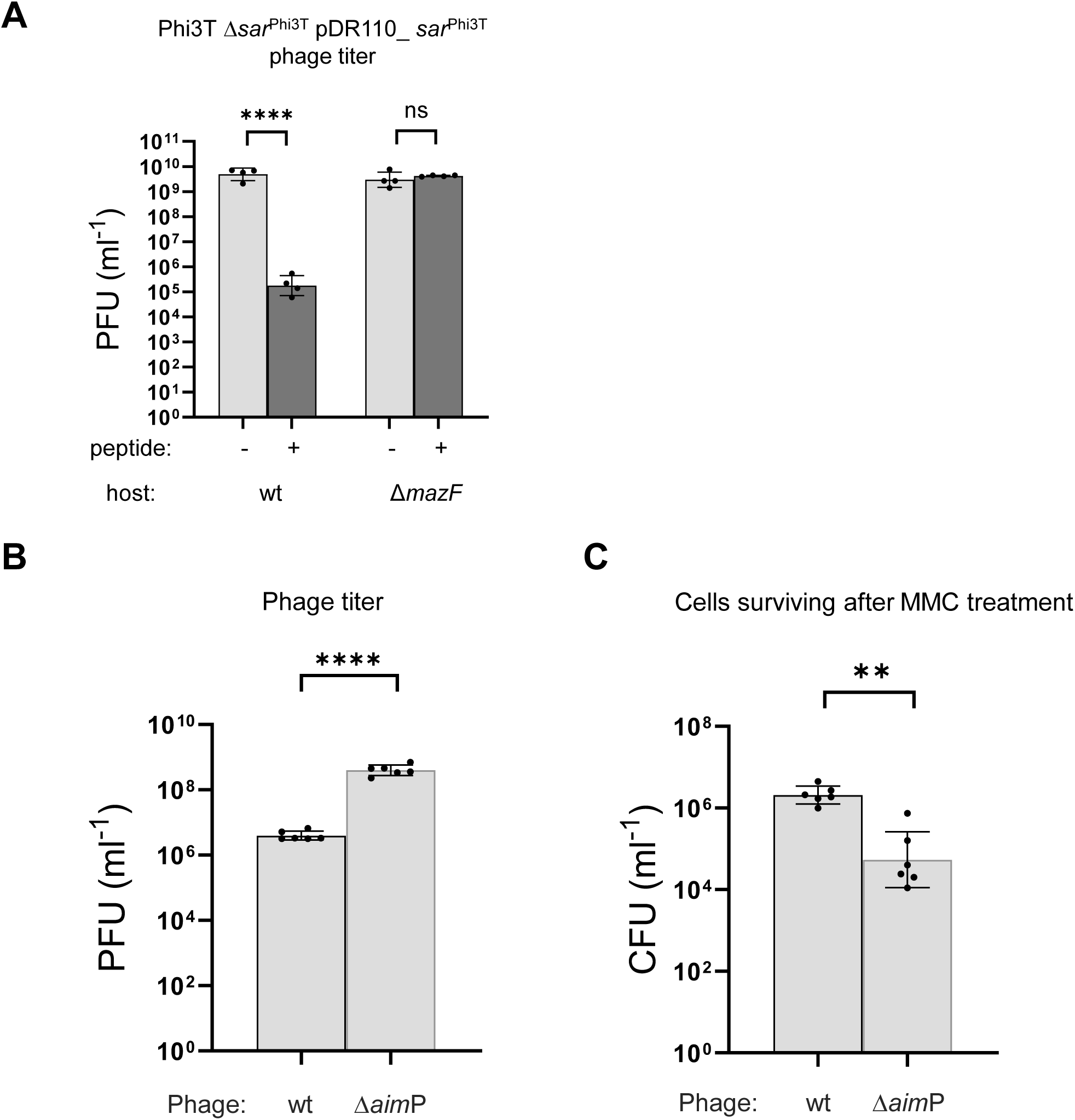
Arbitrium peptide regulation of phage induction happens downstream of Sar^Phi3T^ expression and increases survival of lysogens upon SOS stress. (**A**) wt or Δ*maz*F strains lysogenic with phage Phi3T Δ*sar*^Phi3T^ were complemented with integration vector harbouring an IPTG-inducible Sar^Phi3T^. The phages were induced with IPTG in the presence or absence of added synthetic arbitirum peptide (5μM) as indicated by +/- symbols. The number of resulting phages were quantified using 168 Δ6 as a recipient strain. The results are shown as PFUs ml^-1^, with geometric means and SDs presented (n = 3). (**B**) Strains lysogenic for wt Phi3T phages or ΔaimP (not producing the arbitirum peptide) were induced with MMC. The number of resulting phages were quantified using 168 Δ6 as a recipient strain. The results are shown as PFUs ml^-1^, with geometric means and SDs presented (n = 6). (**C**) The number of bacteria surviving the MMC treatment was quantified and presented as CFUs ml^-1^, with geometric means and SDs marked (n = 6). In all panels, two-tailed t-tests on log_10_ transformed data were performed to compare mean differences. For panel (C) Welch’s correction was applied. p values are indicated above the plots: p < 0.0001 (∗∗∗∗), p < 0.005 (∗∗), ns not significant.

We then repeated the experiment using a Δ*mazF* mutant. As predicted, in the absence of MazF, the addition of the peptide failed to suppress phage induction triggered by Sar^Phi3T^ expression (Figure 6A). This result confirms that full phage induction requires not only the expression of the phage-encoded antirepressor Sar^Phi3T^ but also a permissive signal from the arbitrium system. In other words, the system functions as a “green light” that integrates environmental information to regulate the lytic-lysogenic decision. When the concentration of arbitrium peptide in the culture is high, signalling the presence of many identical phages, it becomes evolutionarily advantageous to suppress further induction. By maintaining the lysogenic state, the host cells (and thereby the resident prophages) are preserved, preventing unnecessary host lysis and helping to protect the phage population as a whole.

To test whether the arbitirum-mediated suppression increases chances of bacterial survival, we MMC induced the strains harbouring lysogens of Phi3T wt or Phi3T Δ*aim*P (not producing the arbitrium peptide) phages. As expected, when the Phi3T phages produced the arbitirum peptide, less phages were released (Figure 6B) and more lysogens survived in the MMC treated culture as compared to Δ*aimP* Phi3T lysogens (Figure 6C).

### SPBeta phage is also induced by a small antirepressor protein regulated by LexA

The *sroA*-*sroF* operon is conserved in genomic structure but not in sequence between the two model arbitrium phages, Phi3T and SPbeta. SroF^Phi3T^ and SroF^SPBeta^ show only 30% amino acid sequence identity, but they both share a conserved tyrosine integrase-like folding. Therefore, we hypothesised that the induction mechanisms would also be conserved. However, a basic BLASTp search could not identify any homologues of Sar^Phi3T^ among the SPBeta phage ORFs. Therefore, we performed a structural search in the PDB database using DALI server.^39^ This search identified, with a high Z score, YopT (72 aa) (PDB: 2DLB^40^) from SPbeta phage as a structural homolog of Sar^Phi3T^. Importantly, *yop*T is also located in the same genomic context (1 ORF downstream of SroF^SPBeta^) as *sar^Phi3T^* in phage Phi3T (Supplementary Figure S5A). Although the function of YopT was unknown and the protein varies from Sar^Phi3T^ in length and sequence (< 20 % sequence identity), both structures are dimers with high similarity (RMSD of 3.5 Å^2^ for the superimposition of the dimers). This suggested a similar function and led us to verify if YopT (hereafter Sar^SPBeta^) in SPBeta phage also is an antirepressor protein.

To test this idea, we repeated the *in vivo* experiments performed for Sar^Phi3T^, but this time with SPBeta phage and Sar^SPBeta^ and obtained similar results as for Sar^Phi3T^ (deletion mutant phenotype - Supplementary Figure S5B-D; SOS-independent phage induction by IPTG-regulated expression of Sar^SPBeta^- Supplementary Figure S6A-C; SOS-box regulation of *sar*^SPBeta^ promoter - Supplementary Figure S7A-E). This confirms that both Sar^Phi3T^ and Sar^SPBeta^ are antirepressors regulated by the SOS response via lexA biding to SOS boxes located upstream of the antirepressor genes. Importantly, the expression of non-cognate antirepressors did not lead to SPBeta or Phi3T phage induction (Figure 3D, Supplementary Figure S6B), showing that there is specificity between these pairs of master repressors and their cognate antirepressor proteins. Our previous results also showed that the repressor proteins SroF of phages Phi3T and SPBeta cannot block infection by the non-cognate phage, as was predicted based on the different DNA motifs recognised by the different SroFs.^25^ Therefore, both the repressors and antirepressors are phage-specific.

### Widespread conservation of the Sar–SroF switch in arbitrium phages

We hypothesised that the novel induction mechanism involving Sar and SroF is not limited to the two prototypical arbitrium phages studied, but is broadly conserved across the arbitrium phage family. To test this, we constructed phylogenetic trees for both Sar and SroF homologues using a previously established database of SPβ-like phages.^38^ Comparison of these trees revealed a strong pattern of co-evolution between the antirepressor proteins and their cognate SroF partners (Supplementary Figure S8, Supplementary Table S5).

To further assess structural conservation, we modelled representative Sar and SroF proteins using AlphaFold3 (Supplementary Figure S9).^41^ The resulting models revealed conserved folding patterns across diverse phages, supporting the functional conservation of this regulatory module. Additionally, analysis of the genomic context confirmed the presence of SOS boxes near the putative *sar*-like genes in all analysed genomes, indicating that LexA-mediated regulation of antirepressor expression is also a conserved feature (Supplementary Table S5).

Altogether, these findings demonstrate that the Sar–SroF switch, controlled by the SOS response and integrated with the arbitrium communication system, represents a broadly conserved strategy used by arbitrium phages to finely tune the lysis–lysogeny decision, an elegant evolutionary solution to optimise survival in dynamic microbial communities.

## Discussion

This work uncovers the sophisticated mechanism of SPBeta-like phages repression and how the prophages regulate their induction in response to host stress. At the heart of this system is SroF, a highly unusual repressor with a structural fold akin to tyrosine recombinases - enzymes typically involved in DNA integration and excision.^23,25^ Remarkably, SroF has lost its catalytic activity and instead its recognition and specificity capabilities have evolved to transform into a dedicated DNA-binding repressor, offering a striking example of functional repurposing.^25^

The crystal structure of the SroF-DNA complex reveals that SroF engages DNA as a monomer, using both CB and CAT domains to form a clamp-like architecture around the DNA helix, with high sequence specificity. This contrasts with typical tyrosine recombinases, which function as dimers or tetramers and exhibit lower sequence readout of its target operators.^36,42^ The absence of the catalytic tyrosine, the high readout capacity of its long (24 bp) target operator sequence and the monomeric DNA binding mode likely reflect SroF’s evolutionary divergence from recombinases toward a specialised role as a transcriptional repressor. While SroF is the first phage repressor known to adopt this fold, other recombinase-like proteins (e.g., from phage P4) have been shown to regulate their own expression, suggesting that the structural framework of recombinases may predispose them for such regulatory adaptations.^43^

Interestingly, SroF also differs from canonical phage repressors such as Lambda CI, which function as dimers and form higher-order oligomers upon DNA binding.^44^ Among phage repressors, the only known functional monomeric counterpart is found in some mycobacteriophages, although these proteins present two HTH domains fused by a helical bridge, thus this monomer recognises the DNA in a manner similar to the lambda CI dimer repressor.^45^ These comparisons highlight SroF as a structurally and functionally unique regulator within the phage world.

Phage induction requires de-repression, typically achieved through two major strategies: RecA-mediated cleavage of repressors (as in Lambda) or repressor displacement by antirepressor proteins.^9,17,46^ SPBeta phages use the latter mechanism, encoding a protein antirepressor, Sar, that binds to SroF and displaces its DNA association. To perform this function and compete with the extensive readout that SroF carries out on its operator, Sar employs a dual strategy. On one hand, it mimics DNA, and on the other, it induces a conformational change in SroF. Sar mimicks electrostatic and structural features of the DNA interface recognised by SroF CAT domain and traps SroF CB domain in a non-functional intermediate conformation between the apo and DNA bound form, thereby abolishing its repressive function. The later mechanism resembles that of the Salmonella phage SPC32H antirepressor Ant,^47^ although Sar additionally occludes the DNA-binding domain itself, representing a dual mode of inactivation.

Sar and related antirepressors are emerging as central regulators of prophage induction. Many are themselves regulated by LexA, linking phage induction to the host SOS response.^16,17,19,20^ This is especially important in SPBeta phages, where SroF is not susceptible to RecA-mediated cleavage. In such systems, antirepressors act as critical molecular switches that translate host stress into prophage induction. However, not all antirepressors follow this logic. Some are not LexA-regulated and instead respond to alternative signals, such as quorum sensing molecules produced by the host. Phages like VP882 and ARM81ld monitor host population density cues via such quorum-sensing-regulated antirepressors, representing an SOS-independent mode of induction.^18,48^ Since VP882 and ARM81ld phages encode repressors that can undergo RecA-mediated cleavage, these systems offer phages the flexibility to tune their lifecycle to both intracellular stress and extracellular ecological conditions.^18,48^

SPBeta-like phages are able to integrate both SOS and arbitrium signalling to control induction.^22^ The arbitrium system coordinates phage decision-making based on the abundance of infected or lysogenic cells present in the environment, while the SOS pathway signals host distress.^21–23^ Importantly, overexpression of the arbitrium receptor AimR alone is insufficient to trigger induction, indicating that both pathways are needed for robust activation.^23^ This dual-input logic makes biological sense: the presence of the arbitrium peptide signals that other phages in the population have already induced or that the majority of surrounding bacteria are already lysogenized by the same phage, thus unsusceptible to further infections. In this context, it may be more advantageous for a prophage to remain latent and help preserve the viability of its lysogen, especially if that prophage can encode genes that enhance host survival.^49^ Thus, integrating stress and quorum-based signals enables SPBeta-like phages to adopt a more nuanced, population-aware strategy - inducing only when the host is damaged and uninfected hosts are likely to be available, while otherwise favouring lysogeny to maintain long-term fitness.

Another key insight from this study is the potential specificity of Sar activity, since between the model SPBeta and Phi3T phages, Sar antirepressors do not induce non-cognate phages. However, cross-inducing antirepressors have been reported in other systems and may enable coordinated induction across co-resident prophages in polylysogenic hosts.^19^ Whether such cross-reactivity exists within the SPBeta family remains an open question and is currently under investigation. If present, these interactions could facilitate phage–phage communication and synchronised induction, reflecting a form of social behaviour that enhances collective fitness. Importantly, these dynamics are not restricted to prophages alone. Both phages and phage satellites have been shown to encode antirepressors that target one another to modulate induction, as exemplified by cases where phage or satellite proteins act as antirepressors of satellite repressors.^50–52^ Importantly, given their small size, low sequence conservation, and structural diversity, antirepressors remain difficult to identify bioinformatically. As a result, the true scale and ecological impact of these proteins is likely underestimated. Our findings expand the known repertoire of antirepressor mechanisms and highlight the importance of structural biology in unravelling their mode of action.

While this study focuses on the antirepressor-mediated inactivation of the master repressor SroF, recent work has identified a second repressor, SroE, encoded by SPBeta-like phages.^25^ SroE is also required for establishing lysogeny, suggesting that it plays a non-redundant role in the regulatory network controlling the lysis-lysogeny switch. How SroE fits into the current model of phage induction remains unclear. One possibility is that SroE may function to stabilise the SroF-DNA complex, reinforcing repression under non-inducing conditions. Another alternative is that it modulates key regulatory proteins involved in the lysis–lysogeny decision-making process, such as AimR, AimX, or Sar, in phages with arbitrium communication system. However, these hypotheses remain untested, and the precise molecular relationship between SroF and SroE is still to be elucidated. The presence of two repressors with potentially complementary roles underscores the complexity of the regulatory circuitry in SPBeta-like phages and points to the existence of additional layers of control that fine-tune the phage lifecycle.

In conclusion, this study uncovers the molecular logic by which SPBeta-like phages regulate their lysis-lysogeny decision, centered on the integrase-like repressor SroF and its dedicated antirepressor Sar. This system exemplifies a sophisticated integration of host stress sensing via the SOS response and phage–phage communication through the arbitrium quorum-sensing pathway. By tightly linking environmental signals to genetic switches, these phages ensure induction occurs only under optimal conditions - when the host is compromised, and new hosts are likely available. This work highlights a previously unappreciated class of repressor–antirepressor pairs and expands our understanding of how temperate phages adapt their regulatory circuits to thrive in complex, competitive microbial communities.

## ACKNOWLEDGEMENTS

We thank the IBV-CSIC Crystallogenesis Facility for protein crystallization screenings. Data collection experiments for the crystallographic structures were carried at XALOC beamline at ALBA Synchrotron with the collaboration of ALBA staff. X-ray diffraction data collection was supported by block allocation group (BAG) ALBA Proposal 2024078509. We acknowledge support from the Scientific Network Conexion Resistencia Antimicrobianos funded by the Consejo Superior de Investigaciones Científicas (CSIC), Spain. This work was supported by grants PID2022-137201NB-I00 from Spanish Government (Ministerio de Ciencia e Innovación), CIPROM/2023/30 from Valencian Government and the European Commission NextGenerationEU fund (EU 2020/2094), through CSIC’s Global Health Platform (PTI Salud Global) to A.M; and grants MR/X020223/1, MR/M003876/1, MR/V000772/1 and MR/S00940X/1 from the Medical Research Council (UK), BB/V002376/1 and BB/V009583/1 from the Biotechnology and Biological Sciences Research Council (BBSRC, UK), and EP/X026671/1 from the Engineering and Physical Sciences Research Council (EPSRC, UK) to J.R.P. A.M, A.E and J.R.P are funded by European Research Council Grant 101118890 (TalkingPhages).

## AUTHOR CONTRIBUTIONS

A.M. and J.R.P. conceived the study; C.C., S.Z.-C., J.M.-B., Y.L., D.S., T.B., and S.O.B. conducted experiments; C.C., S.Z.-C., J.M.-B., Y.L., D.S., T.B., S.O.B., A.E., M., and J.R.P. analyzed the data; C.C., S.Z.-C., A.M. and J.R.P. wrote the manuscript, with inputs from the rest of the authors.

## DECLARATION OF INTEREST

The authors declare no competing interests.

## MATERIAL AND METHODS

### Bacterial strains and growth conditions

Strains used in this study are listed in Supplementary Table S6. *E. coli* and *B. subtilis* strains were grown at 37 °C, shaking at 120 rpm in liquid broth (LB Lennox or Miller, respectively) or plated onto corresponding LB media bacteriological agar plates (1.5% w/v). In all liquid cultures Miller LB was supplemented with 0.1 mM MnCl_2_ and 5 mM MgCl_2_. When indicated, antibiotics were added at the following concentrations: erythromycin (1 μg ml^-1^), kanamycin (10 μg ml^-1^), ampicillin (100 μg ml^-1^), spectinomycin (100 μg ml^-1^), chloramphenicol (5 ug ml^-^ 1). All reagents are listed in Supplementary Table S7.

### Deletion mutant construction

Phages and phi3T and SPβ with kanamycin markers were used in this study.^23^ To generate the deletion mutants of *sar*^Phi3T^ and *sar*^SPBeta^ genes, we followed the protocol from.^53^ We first generated overlapping PCRs containing 1kb flanking regions of target genes and an erythromycin cassette flanked by lox sites. The PCRs were transformed into Δ6 lysogen strains. Competent *B. subtilis* cells were prepared as previously described.^54^ Insertion of the erythromycin cassette was confirmed by Sanger sequencing. The cassette was then removed by transforming pDR244 (selection for spectinomycin resistance) at 30 °C to allow for Cre/lox-mediated removal of the cassette. Transformant colonies were then restreaked onto LB plates and incubated overnight at 42 °C to remove the temperature-sensitive plasmid. The colonies were screened for the loss of plasmid and erythromycin cassette (spectinomycin and erythromycin sensitive), and the deletion was confirmed by PCR and Sanger sequencing.

### Plasmid construction

All plasmids and primers are listed in Supplementary Table S4. For cloning genes into vectors, the target genes were amplified by PCR with either (1) restriction enzyme sites introduced into overhangs or (2) overlapping sequences enabling Gibson assembly (listed in Supplementary Table S6). The digested fragments were ligated with plasmids linearised using the same enzymes or Gibson assembly was performed using NEBuilder HiFi DNA Assembly master mix (New England Biolabs), as specified in Supplementary Table S6. Next the constructs were transformed into *E. coli* DH5alpha, DH12 or BL21. Sequencing was used to confirm the correct constructs. In case of integrative expression vectors, the constructs were then transformed into *B. subtilis* strains with selection for the corresponding antibiotic (spectinomycin for pDR110 derivatives, erythromycin for pDG1663_GFP, pAX01 and pDG1664 derivatives, and chloramphenicol for pAEC310 derivatives). The double recombination of the integration vectors was confirmed by starch tests (pDR110 derivatives), sensitivity to Spectinomycin (pDR1664 derivatives), or PCRs of flanking regions (pDG1663_GFF, pAX01 and pAEC310 derivatives). The integration sites, antibiotic resistance and inducible promoters of plasmids are listed in Supplementary Table S6.

Plasmid pDG1663_GFP was constructed from pDG1663+ by replacing the B-galactosidase gene with sfGFP. First, the pDG1663 plasmid was digested with BamHI and EcoRV. The *sfGFP* gene was PCR amplified from plasmid pAND101 and joined with an overlapping PCR of the flanking thrC fragment from plasmid pDG1663. The PCR was digested with BamHI and EcoRV, ligated with the linearised pDG1663 and transformed into *E. coli*.

### Phage induction assay

For induction, overnight *B. subtilis* Δ6 lysogen cultures were diluted to OD_600nm_ = 0.05 in LB and grown at 37 °C with 120 rpm shaking until OD_600_ = 0.2. This step was repeated twice to ensure cells were in exponential growth. When indicated, Mitomycin C (MMC) was added to final concentration of 0.5 μg ml^-1^, and/or IPTG to final concentration of 1 mM (complementation experiments). In induction experiments involving synthetic peptide, the Phi3T 6-aa mature arbitrium peptide (SAIGRA; ThermoFisher Scientific; >95% purity, desalted) was added to final concentration of 5 μM simultaneously with MMC. The induced cultures were then incubated at 30 °C with 80 rpm shaking for 4 h followed by overnight incubation without shaking at room temperature. Cultures were then filtered (0.2 μm) and lysates were stored at 4 °C until use.

### Titration assays

A. *subtilis* Δ6 was the recipient for quantifying phages after induction. Overnight cultures of *B. subitilis* Δ6 were diluted to OD_600nm_ 0.05 in LB media and shaken at 120 rpm, 37 °C until an OD_600nm_ value of 0.2. 100 μl of bacterial culture were then mixed with 100 μl of serial dilutions of the lysates in phage buffer (1 mM NaCl, 0.05 M Tris pH 7.8, 0.1 mM MnCl_2_, 5 mM MgCl_2_). After 10 minutes at room temperature, the mixture was mixed with 3 mL of 0.5 % top agar and immediately plated onto LB plates. The top agar and LB plates were both supplemented with 0.1 mM MnCl_2_ and 5 mM MgCl_2_. The top agar was left to dry for 20 minutes, then incubated at 37 °C degrees overnight. Plaques were counted the next day.

### Lysogeny assays

Lysogens were quantified by growing *B. subitilis* Δ6 to OD_600nm_ value of 0.2 in LB. Lysates were serially diluted in phage buffer and 100 μl was added to 1 ml of the recipient bacteria. After incubating at 37 °C for 30 min to allow the phage to infect bacteria, the mixtures were centrifuged at 6600 rpm for 1 min. The pellets were resuspended in 100 μl of LB and plated onto selective antibiotic LB agar plates. Plates were incubated overnight at 37 °C. The number of colonies forming units (CFU) was counted.

### Phage induction and protein expression growth curves

Overnight cultures of *B. subitilis* strains Δ6 harbouring the integration vectors or harboring prophages and integration vectors as indicated were diluted to OD_600nm_ = 0.05 in LB and shaken at 120 rpm, 37 °C until OD_600nm_ = 0.2. When indicated, MMC was added to final concentration of 0.5 μg ml^-1^, and/or IPTG to final concentration of 1mM and 200 μl of culture were transferred into 96-well plates Cell growth and lysis was then followed by OD_600nm_ measurements in a 96-well plate reader (BMG Labtech FLUOstar Omega) for 6 hours.

### Phage infection growth curves

To test the behavior of phages in infection, overnight cultures of *B. subitilis* Δ6 were diluted to OD_600nm_ = 0.05 and shaken at 120 rpm, 37 °C until OD_600nm_ = 0.2. Phages were then added to the culture at an MOI of 0.1 and OD_600nm_ measurements were taken in a 96-well plate reader (BMG Labtech FLUOstar Omega) for 6 hours.

### Evolution experiments

To evolve phages that cannot be induced by *sar*^Phi3T^ expression, we used a *B. subitilis* Δ6 Phi3T Δ*sar*^Phi3T^ lysogen harbouring two copies of *sar*^Phi3T^ (under xylose and IPTG inducible promoters). Overnight lysogen cultures were diluted to OD_600nm_ = 0.05 in LB and grown at 37°C with 120 rpm shaking until OD_600_ = 0.2. At that point IPTG and xylose were added to final concentrations of 1mM and 2% (w/v), respectively. The induced cultures were then incubated at 30 °C with 80 rpm shaking for 4 h. 100 μl of the samples were then plated on agar plates supplemented with kanamycin. Surviving single colonies were re-streaked and tested again for induction with xylose and IPTG. The cultures that showed no or bad lyses were filtered for phages. Next, the lysates of putatively evolved phages were used to form lysogens in a new recipient strain harbouring a copy of IPTG inducible *sar*^Phi3T^ and tested for lysis upon adding IPTG. This allowed us to exclude instances when bad induction was due to chromosomal mutations. Candidate lysogens were then sequenced (Nanopore sequencing, Plasmidsaurus).

### MMC survival experiment

This experiment was performed in minimal media (10.7 g L^-1^ K_2_HPO_4_, 6 g L^-1^ mM KH_2_PO_4_, 0.85 g L^-1^ Na_3_Citrate.2H_2_O, 2% w/v Glucose, 0.05 mg mL^-1^ L-tryptophan, 0.011 mg mL^-1^ Ferric ammonium citrate, 2.5 mg mL^-1^ L-aspartic acid, 3 mM MgSO_4_, based on ^55^), where the difference in phage titer between wt and Δ*aim*P phage induction was reported to be more pronounced.^22^ Overnight cultures of *B. subitilis* Δ6 Phi3T lysogens were diluted to OD_600nm_ value of 0.05 in minimal media and then shaken at 37°C, 120rpm until an OD_600nm_ value of 0.2. MMC was added to a final concentration of 0.5 μg ml^-1^ when indicated. Cultures were shaken at 30 °C, 80 rpm for 12 hours. Next, cultures with MMC added underwent two rounds of washing with phosphate-buffered saline (PBS) solution to remove remaining MMC. To wash the cells, 1mL of culture was pelleted (x8000 g, 5 min), 950 μl of supernatant was removed, followed by resuspension in 950 μl PBS. Cultures were then serially diluted in PBS and 100 μl was spread on LB plates containing kanamycin. The plates were incubated at 37 °C overnight and colonies were counted the next day. 1 mL of the cultures was also extracted and 0.22 μm filtered for phage titration. 2 colonies were also picked for each induction to verify that the colonies that grew were indeed lysogens containing viable prophages. Cultures derived from these colonies were MMC induced and the titers counted.

### GFP and YFP reporter assays

Overnight cultures of the *B. subitilis* strains harboring reporter vectors as indicated were diluted to an OD_600nm_ value of 0.05 in LB. The cultures were then shaken at 37 °C, 120 rpm to an OD_600nm_ value of 0.2, after which they were diluted again in minimal media to OD_600nm_ = 0.05. At this point, for SroF^Phi3T^-Sar^Phi3T^ YFP reporter experiments, IPTG and xylose were added to final concentrations of 1mM and 2% (w/v) respectively to all strains to enable the expression of *sar*^Phi3T^ (IPTG) and *sroF*^Phi3T^ (xylose). The cultures were then shaken at 37 °C, 120 rpm to an OD_600nm_ value of 0.2. 200 µL of the cultures was transferred to a 96-well plate. At this point, for GFP reporter experiments (for both Phi3T and SPBeta *sar* promoters) MMC was added to a final concentration of 0.5 μg ml^-1^ as indicated. OD_600nm_ and fluorescence were then measured every 15 min for 6 h, using 435/535nm and 485/535nm excitation/emission filters for GFP and YFP reporters respectively. In case of the GFP reporters the host strain was *B. subitilis* Δ6. In case of YFP reporters for Phi3T phage constructs the host strain was Δ6s - a chloramphenicol sensitive derivative of *B. subitilis* Δ6, to enable selecting for transformation of the YFP reporter plasmid encoding a chloramphenicol marker.

For SroF^SPBeta^-Sar^SPBeta^ YFP reporters PY79 Δxpf was used as a host in strains AES9539, AES9826 and AES9846. They were grown overnight in LB at 37° with shaking at 220 RPM then diluted by a factor of 1:100 into fresh LB media. Upon reaching OD_600nm_ = 0.3 cultures were supplemented with blank (AES9539), 1mM IPTG (AES9826), or 1mM IPTG and 2% xylose (AES9846) and grown for an additional 90 minutes, after which YFP levels were measured via flow-cytometry. Flow cytometry was performed to quantify gene expression at the single-cell level, using a Beckman-Coulter Cytoflex flow-cytometer equipped with four lasers (405 nm, 488 nm co-linear with 561 nm, 638 nm). The emission filters used were: YFP – 525/40. Median YFP over >20,000 cells were measured in each experiment and the experiment was repeated three times.

### Recombinant protein expression and purification

Constructs with *sroF^Phi3T^* and *sar^Phi3T^* genes cloned in pLIC-SGC1 plasmid were transformed into an *E. coli* strain BL21 (DE3) RIL (Agilent) for protein overexpression, that was performed as described in ^26^. Briefly, for each construct a single colony carrying the expression plasmid was grown overnight at 37 °C in 50 ml of LB medium supplemented with 50 µl of 100 μg ml^-1^ ampicillin solution and 50 µl of 33 μg ml^-1^ chloramphenicol solution. The culture was used to inoculate 1 L of LB medium, containing ampicillin and chloramphenicol, and was grown at 37°C until an OD_600nm_ reached a value between 0.6 - 0.8. At that point, the temperature was set to 20 °C and protein expression was induced with a final concentration of 1 mM of IPTG. The culture was incubated at 20 °C for additional 16 hours. Cells were harvested by centrifugation, washed with PBS and frozen at −20 °C until used.

Purification of both proteins was performed as follows. First, pellets were resuspended in lysis buffer (50 mM Tris pH 8, 500 mM NaCl) and sonicated 10 minutes in ice, with pulses having 1 second on 1 second off to avoid heating of the sample. Sonicated sample was centrifuged at 10.000 *g* for 1 h to separate cell debris. The supernatant was filtered using a 0.45 nm filter and loaded into a 1 ml His-Trap (Cytiva) column previously equilibrated with buffer A (50 mM Tris pH 8, 250 mM NaCl). After a washing step with buffer A, proteins were stepwise eluted with buffer B (buffer A with 500 mM imidazole). Fractions containing the purest protein were pooled and digested, if needed, with TEV protease (50:1 molar ratio protein:TEV). The sample was concentrated and loaded into a Superdex 75 increase 10/300 (Cytiva) gel filtration column equilibrated in buffer A. The purest fractions judged by SDS-PAGE were pooled, concentrated and stored at −80 °C until use.

For structural studies, pure SroF^Phi3T^ and Sar^Phi3T^ were mixed, and the complex was purified by molecular exclusion chromatography using a Superdex 200 increase 10/300 (Cytiva).

Since in both, Genbank and Uniprot the Sar^Phi3T^ sequence (Phi3T_99) was annotated as a 110 amino acids protein, we used such sequence to clone and produce recombinant Sar^Phi3T^. However, upon solve SroF^Phi3T^-Sar^Phi3T^crystal structure (see below), the first 10 amino acids of Sar^Phi3T^ were not visible in any of the two molecules of the dimeric structure. That lead us to analyse protein annotation, discovering that in the DNA sequence corresponding to those “extra” 10 amino acids one can find a putative ribosome binding site. Based on this finding and in the crystal structure we conclude that the sequence is wrongly annotated and that the protein lacks those 10 first amino acids. We cloned and produced the corresponding Sar variant, following the same protocol and producing a protein that behaves identically to 110 amino acids long version. Therefore, all other experiments, including all *in vivo* experiments, were performed using constructs corresponding to the shorter annotation.

### Protein crystallisation

Crystals of the SroF^Phi3T^-DNA complex were generated by co-crystallization after mixing SroF^Phi3T^ at 20 mg ml^-1^ with the duplex DNA at 1:1 molar ratio. Crystal screens were set up by mixing 0.3 µl of sample and 0.3 µl reservoir solution at 21 °C, using the sitting drop vapor diffusion method. Crystals grew in JBS I C8 (Jena Bioscience) condition: 25 % PEG 4000, 0.1 M Tris pH 8.5 and 200 mM CaCl_2_.

SroF^Phi3T^-Sar^Phi3T^ crystals were grown also at 21 °C using the sitting drop method by mixing 0.3 µl protein complex solution at 13 mg ml^-1^ and 0.3 µl reservoir solution of Wizard HT crystal screen (Hampton) F12 condition: 0.1 M Imidazole pH 8, 30 % PEG 8000 and 200 mM NaCl.

### Data collection, structure solution and refinement

Prior to data collection, crystals were flash frozen in liquid nitrogen. For SroF^Phi3T-^Sar^Phi3T^ crystals, the corresponding crystallization solution was supplemented with 10% glycerol as cryoprotectant. All the diffraction experiments were carried out at 100 K.

Both, SroF^Phi3T^-DNA and SroF^Phi3T-^Sar^Phi3T^ (PDB codes 9RD5 and 9RGL respectively) diffraction data were collected at Xaloc beamline, ALBA synchrotron in Barcelona, using a wavelength of 0.9793 Å in both cases. For SroF^Phi3T^-DNA, data reduction and scaling was carried out in CCP4i^56^, whereas SroF^Phi3T-^Sar^Phi3T^ diffraction data reduction was carried out with XDS^57^ and scaling was performed in CCP4i^56^, at 3.05 Å resolution due to the highly anisotropic nature of the diffraction. For SroF^Phi3T^–DNA phases were calculated using separated domains of SroF^SPBeta^ as templates and Phaser^51^as implemented in Phenix.^58^ SroF^Phi3T-^Sar^Phi3T^ initial phases were determined using ModelCraft^59^ and the separated domains of SroF^Phi3T^ from the DNA bound structure as templates. Followed phase determination, in both cases model refinement was carried out combining cycles of manual building with Coot^60^ and computational refinement using phenix.refine^58^.

### Pull-down assays

Overnight cultures of *B. subtilis* Δ6SroF^Phi3T^(FLAGx3) lysogenized with Phi3T phage were diluted to OD_600nm_ = 0.05 in 50 ml LB (Miller) and incubated at 37 °C 230 rpm. ^25^ When the OD_600nm_ reached around 0.4-0.6, 1 mM IPTG was added. After 3 hours at same growth conditions, cells were harvested and resuspended in PBS. Cells were incubated with 2 % formaldehyde for 20 min at room temperature. After the addition of glycine to stop the crosslinking reaction, cells were washed with TBS and kept in −20 °C at least 24 h. Lysis of the cells was performed using BugBuster (Milipore) combined with sonication in BioruptorPICO (Diagenode). Anti-DYKDDDK magnetic agarose (ThermoScientific) was used to immunoprecipitate SroF^Phi3T^(FLAGx3). After washing and elution, samples were incubated at 65 °C overnight with shaking to 800 rpm. Samples were concentrated using the concentrator Vacufageplus (Eppendorf) and proteins were identified by LCMS/MS.

### EMSA competition assay

SroF^Phi3T^ DNA binding fragment (122 pM) was incubated with SroF^Phi3T^ (37.5 µM) for 5 min at 4 °C. After that time, increasing amounts of Sar^Phi3T^ were added into the different tubes so SroF^Phi3T^ – Sar^Phi3T^ 1:0.25, 1:0.5, 1:0,75, 1:1 and 1:2 ratios were tested to see the effect of Sar^Phi3T^ on SroF^Phi3T^ DNA binding. DNA binding fragment alone, DNA incubated with Sar^Phi3T^ and DNA incubated with SroF^Phi3T^ were used as controls. After Sar^Phi3T^ addition, samples were incubated 10 min at 4 °C prior to be loaded into an acrylamide gel. DNA binding fragment alone, DNA incubated with Sar^Phi3T^ and DNA incubated with SroF^Phi3T^ were used as controls. EMSA was performed at 4 °C, running the gel at 90 V during 70 min. For DNA visualization, the acrylamide gel was incubated in TBE 1X solution with 0.025 % of GreenSafe Premium solution (NZYtech).

### Binding kinetics analysis

Binding affinities were measured at room temperature using the Octet Red96e system (Sartorius). All the proteins were diluted in binding buffer (25 mM Tris pH 8, 150 mM NaCl, 3 mM Imidazole and 0.005% tween 20). Measurements were carried out by immobilizing SroF^Phi3T^ at 0.2 mg ml^-1^ on Ni-NTA biosensors (Sartorius) through its N-terminal 6 His-tag. After generating a baseline with binding buffer, association and dissociation were monitored for 200 seconds each, using a Sar^Phi3T^ concentration range from 500 to 7.813 nM. Binding parameters were calculated using Octet Analysis Studio Software (Sartorius) and a bivalent analyte model.

### Phylogeny construction

We utilised our previously established database of SPBeta-like phages, comprising 321 SPβ- like genomes identified from Bacillus rRNA group 1 strains.^38^ In this study, we specifically searched for homologs of two proteins, SroF and Sar, across all predicted open reading frames (ORFs) in the database. Protein homology searches were conducted using DIAMOND v2.1 with default settings against the annotated ORF sets.^61^ ORFs matching SroF and Sar were identified based on sequence similarity and manually verified when necessary to ensure correct annotation. Notably, Sar^Phi3T^and Sar^SPBeta^ were not found to be homologous and were clustered into two separated Diamond clusters, which were joined in order to make the single phylogenetic tree. Multiple sequence alignments were performed using MAFFT v7.525^62^, and phylogenetic trees were constructed with FastTree v2.1.^63^ Visualization of the resulting trees was carried out using iTOL v7.1.^64^ Structural models for the different SroF and Sar repressentatives were performed with Alphafold 3^41^.

### Identification of LexA sites in SPBeta-like phages database

LexA binding sites were identified using a custom MATLAB script that scanned sequences with the LexA position-specific scoring matrix (PSSM) obtained from the Database of Transcriptional Regulation in *Bacillus subtilis* (DBTBS).^65^ Specifically, the regions from 200 bp upstream to 100 bp downstream of the *sar* start codon were searched. Sites with a PSSM score greater than 8.0 were considered putative LexA binding sites.

### Statistical Analysis

Statistical analyses were performed as indicated in the figure legends using GraphPad Prism 9 software. The p-values represented in each figure are shown in the figure legends. In all figure descriptions the number n represents biologically independent experiment results.

## Notes

### Competing Interest Statement

The authors have declared no competing interest.

