## Supplementary material for "An unprecedented DNA recognition–mimicry switch governs induction in arbitrium phages": Supplmentary Figures S1-S9: Supplementary_Figures.pdf

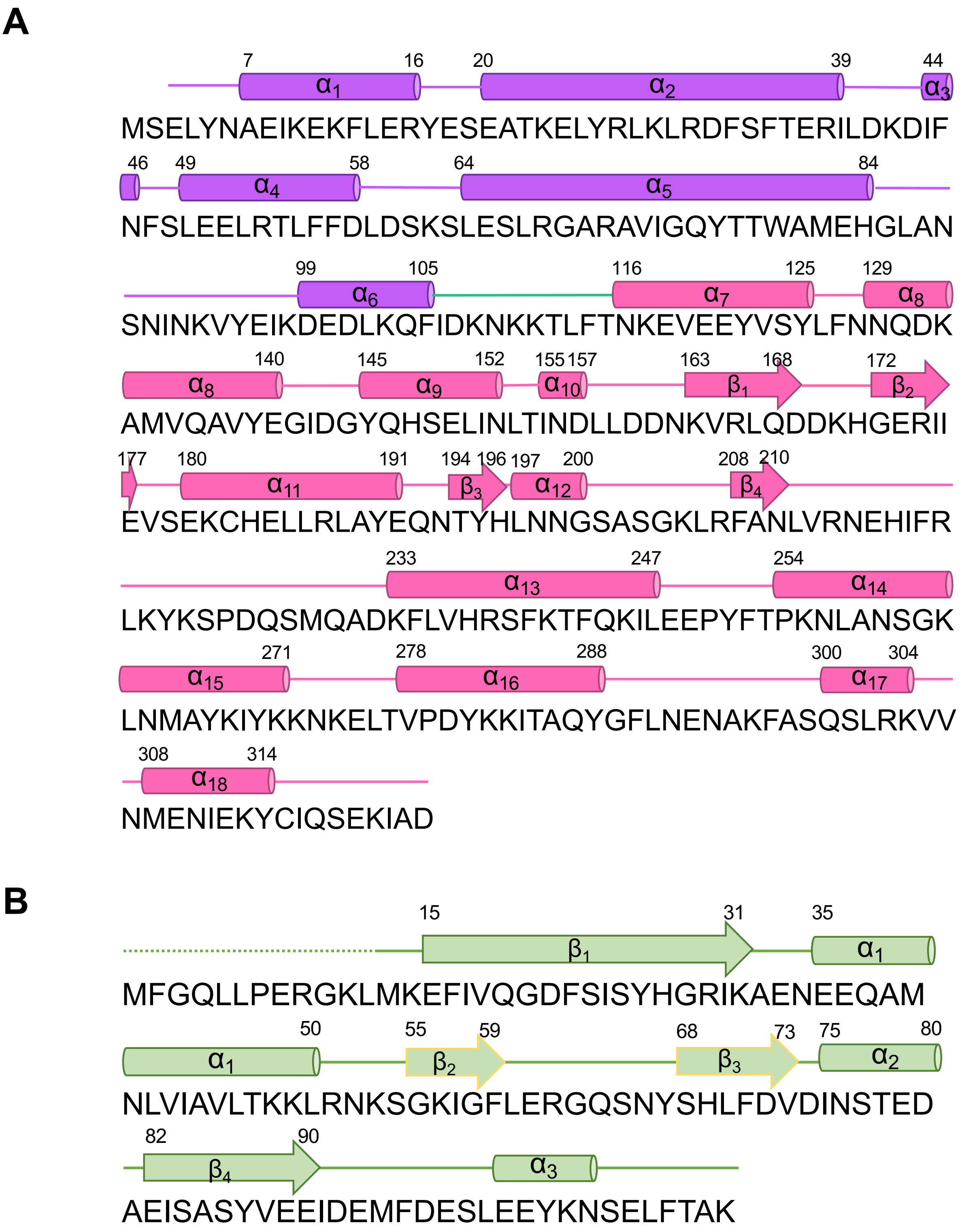


**Supplementary Figure S1. Topology diagrams for SroF$^{Phi3T}$ and Sar$^{Phi3T}$** (**A**) Topology diagram of SroF$^{Phi3T}$. Secondary structure elements, indicated on top of the sequence, are coloured following the same colour pattern as in Figure 1A. (**B**) Topology diagram of Sar$^{Phi3T}$. Secondary structure elements are indicated on top of the sequence. Dotted line indicates non-visible parts in the crystal structure.

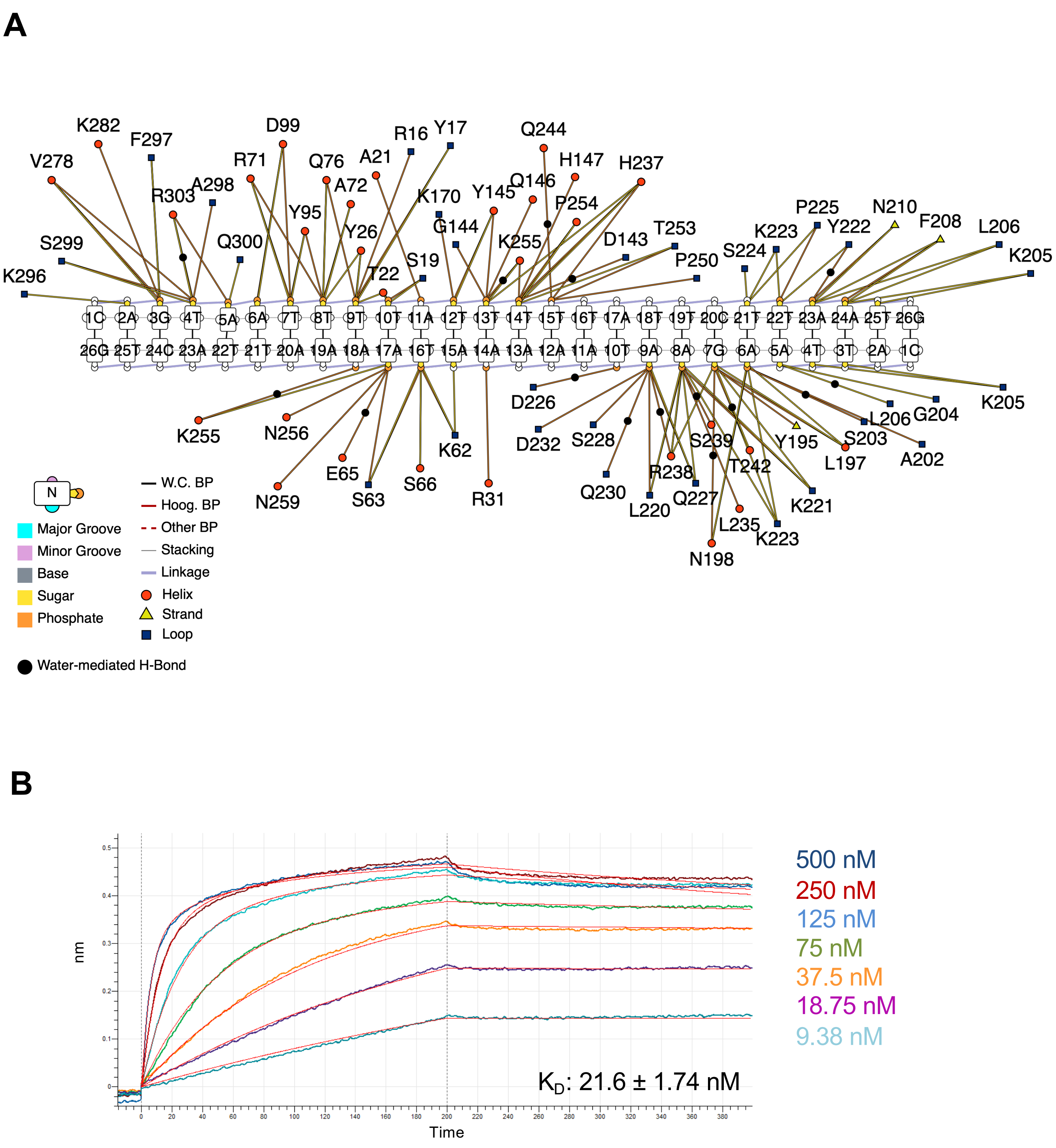


**Supplementary Figure S2. Interactions of SroF$^{Phi3T}$ with DNA and Sar$^{Phi3T}$**
(**A**) Contacts established by SroF$^{Phi3T}$ residues with DNA Backbone in SroF$^{Phi3T}$-DNA structure. Legend indicates the nature of the contact and the secondary structure element where the contacting residue is located. (**B**) Biolayer interferometry measurements of SroF$^{Phi3T}$-Sar$^{Phi3T}$ association and dissociation. Each concentration (from 500 to 9.38 nM) is depicted in a different colour and adjusted model lines are depicted in red. Calculated binding affinity is indicated in the figure.

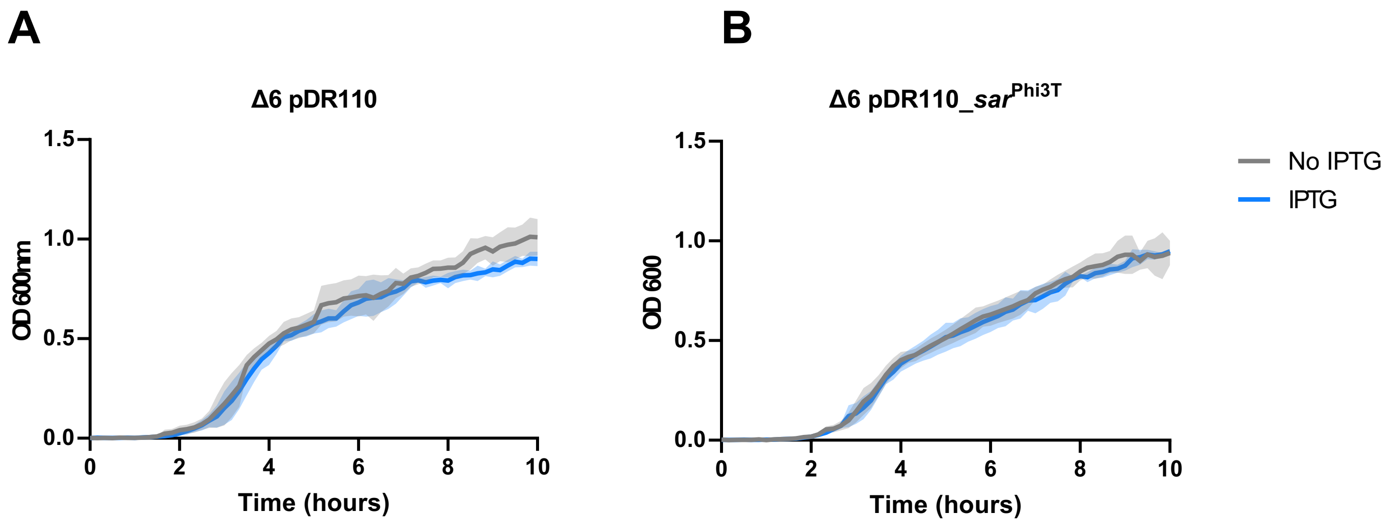


**Supplementary Figure S3 Cell lysis caused by Sar$^{Phi3T}$ expression requires the presence of the phage.** 168 Δ6 strain were complemented with either empty integration vector (amyE::Pspank-) (**A**) or with IPTG-inducible Sar$^{Phi3T}$ (**B**). The $OD_{600}$ readings were measured in absence (grey) or presence of IPTG (blue). Means and SD's are presented (n = 3).

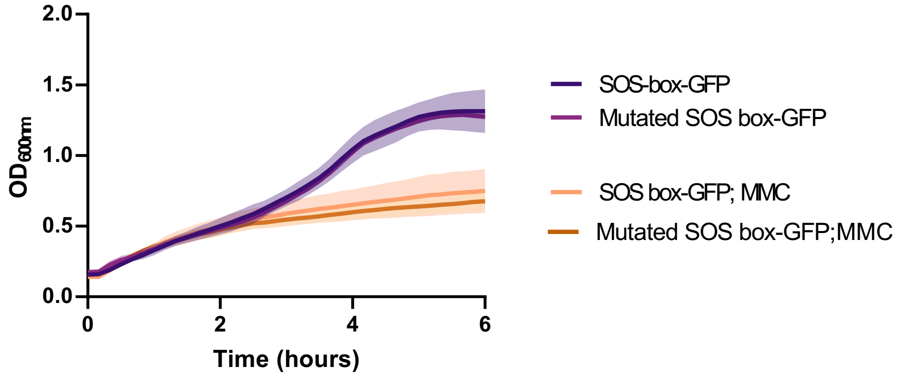


**Supplementary Figure S4 Growth curves of reporter strains in absence or presence of MMC**, corresponding to GFP measurements in Figure 5, panels B-C The $OD_{600}$ readings were measured and means and SD's are presented (n = 3).

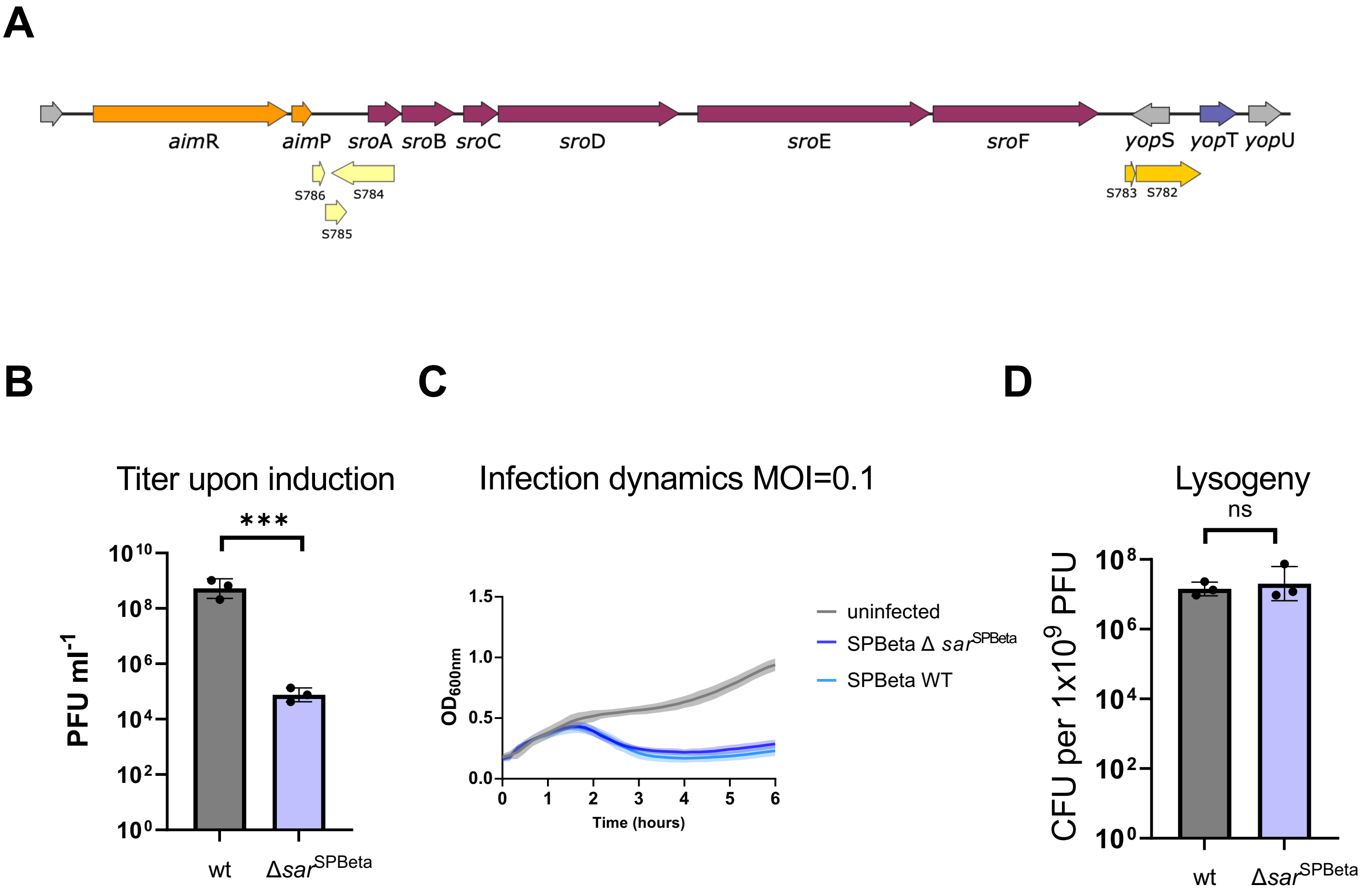


**Supplementry Figure S5. SPBeta phage Sar$^{SPBeta}$ (YopT) characterization. *sar*$^{SPBeta}$ deletion leads to similar phenotypes as deletion of *sar*$^{Phi3T}$ in phage Phi3T. (A)** Schematic overview of the arbitrium operon (orange), 6-gene operon (pink) and *sar*$^{SPBeta}$ (purple) region of phage SPBeta. Putative sRNAs are marked in yellow. (**B**) Strains lysogenic with phages SPBeta wt or SPBeta Δ*sar*$^{SPBeta}$ were MMC induced. Resulting phages were quantified using 168 Δ6 as a recipient strain. (**C**) Strain 168 Δ6 was infected with phages SPBeta wt or SPBeta Δ*sar*$^{SPBeta}$ at MOI=0.1 and infection dynamics were followed by measuring $OD_{600}$ readings. (**D**) The number of lysogens resulting from an infection of 168 Δ6 by SPBeta wt or SPBeta Δ*sar*$^{SPBeta}$ phages. The results are shown in panel (**B**) as PFUs ml$^{-1}$ and in panel (**D**) as CFUs ml$^{-1}$ normalised by PFUs ml$^{-1}$ and represented as the CFU of an average phage titer (1×109 PFU ml$^{-1}$). Data represents geometric mean and SD (n=3) in panels (**B**) and (**D**), and the mean and SD (n=3) in panel (**C**). Two-tailed t-tests were performed on $\log_{10}$ transformed data to compare mean differences. p-values are indicated above each comparison: ***$p < 0.001$; ns – not significant.

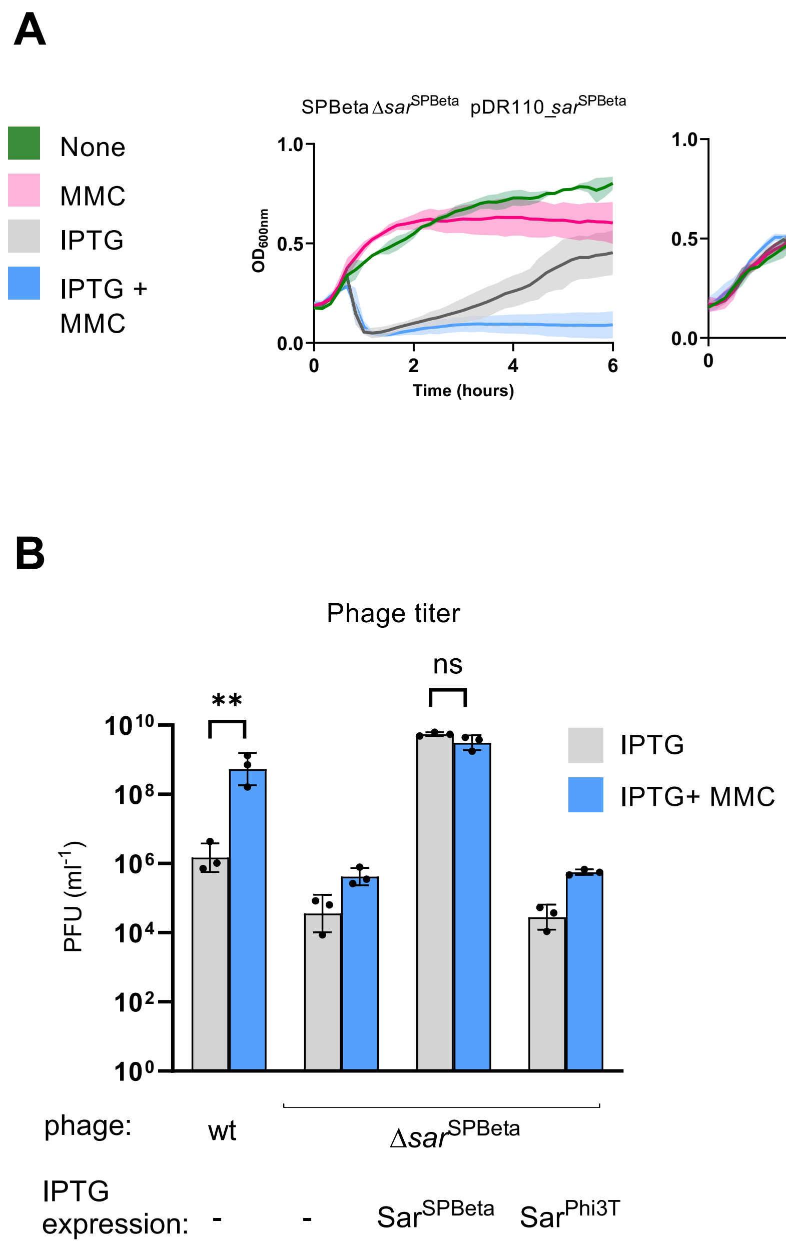


**Supplementray Figure S6. Sar$^{SPBeta}$ expression leads to SPBeta phage induction. (A)** Induction growth curves. Strains lysogenic with phages SPBeta wt or SPBeta Δ*sar*$^{SPBeta}$ were complemented with either empty integration vector (amyE::Pspank-) or with IPTG-inducible *sar*$^{SPBeta}$ or *sar*$^{Phi3T}$. IPTG (gray), MMC (pink) or both IPTG and MMC (blue) were added to the cultures and cell lysis was followed by measuring $OD_{600}$ readings. (**B**) The number of resulting phages upon addition of IPTG (gray) or IPTG and MMC (blue) was quantified using 168 Δ6 as a recipient strain. The results are shown as PFUs $ml^{-1}$, with geometric means and SDs presented (n = 3). A two-tailed t-test on $log_{10}$ transformed data was performed to compare mean differences. (**C**) Sar$^{SPBeta}$ interferes with SroF$^{SPBeta}$ repression of YFP transcriptional reporter constructs. Expression of YFP from yosXY promoter (containing a SPBRE box recognized by SroF$^{SPBeta}$) was measured in the presence or absence (as indicated by the +/- symbols under the x-axis) of xylose-inducible *sro*F$^{SPBeta}$ and/or IPTG-inducible *sar*$^{SPBeta}$. Arbitrary fluorescence unit measurements readings are shown, with means and SDs presented (n = 3). A two-tailed t-test was performed to compare mean differences. ****$p < 0.0001$; **$p < 0.005$; ns – not significant.

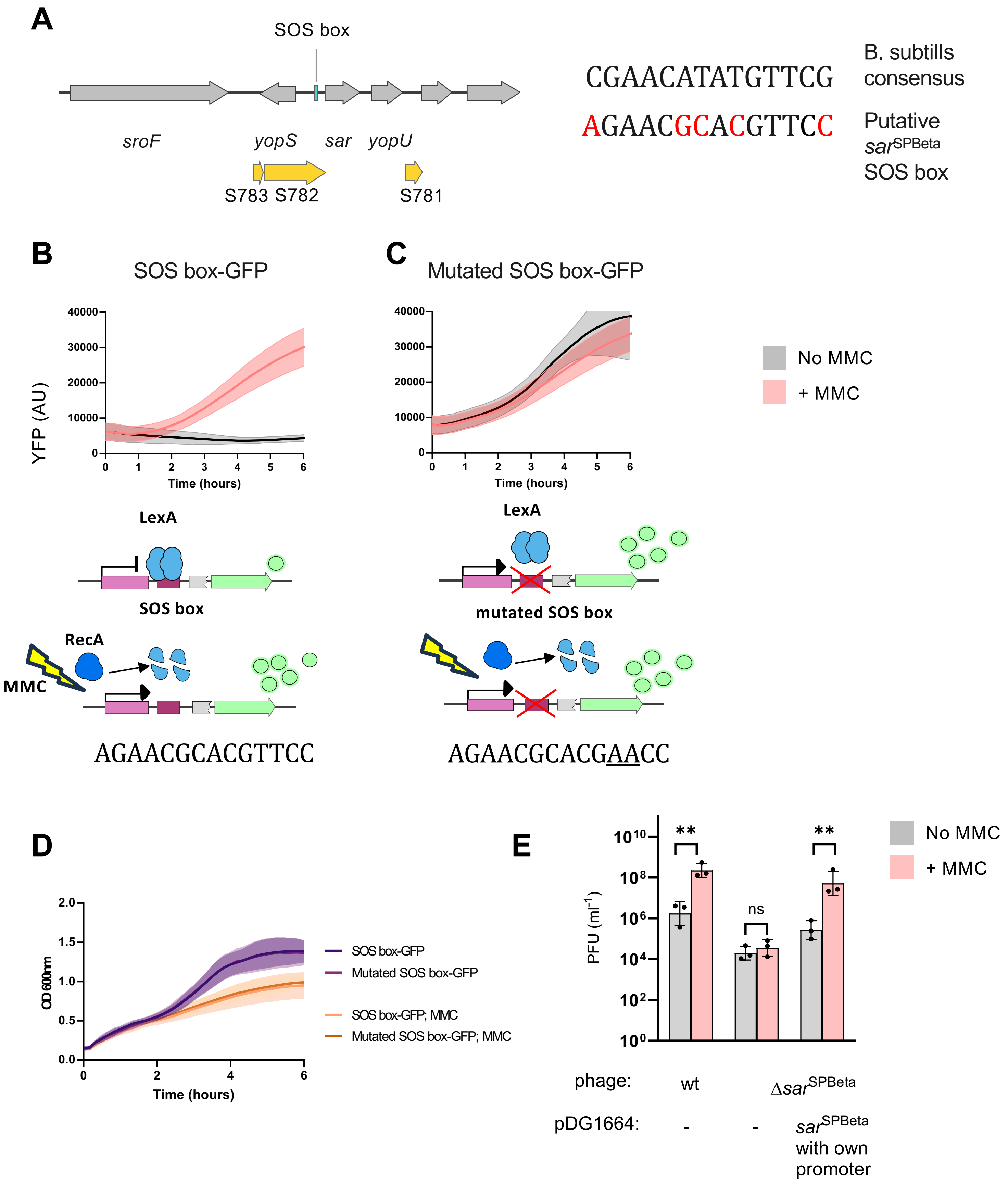


**Supplementary Figure S7. Sar$^{SPBeta}$ (YopT) is regulated by SOS response via LexA-binding.**

**Supplementary Figure S7. Sar$^{SPBeta}$ (YopT) is regulated by SOS response via LexA-binding.** (**A**) Schematic overview of SOS box location upstream of *sar*$^{SPBeta}$ gene. Comparison of consensus sequence for *B.subtilis* SOS-boxes and the putative SOS box upstream of *sar*$^{SPBeta}$, with nucleotide differences marked in red. **(B-C)** Expression of GFP from transcriptional fusion with yopT promoter was measured (arbitrary fluorescence units), with means and SDs presented (n = 3). The expression was measured either in absence (grey) or presence of MMC (pink). The constructs contained either an intact SOS box (**B**) or a version with a mutation in the conserved positions (underlined) (**C**). A schematic of constructs is presented: GFP (green), LexA (blue), RecA (dark blue) *sar*$^{SPBeta}$ predicted promoter (pink), SOS box (dark pink). Due to the reduced growth of cells when treated with MMC (**D**), the fluorescence values were not normalised by OD measurements; however, within a treatment group the strains had similar growth rates, as seen in panel. (**D)** Growth curves of reporter strains in absence or presence of MMC, corresponding to GFP measurements in panels (**B-C**). The $OD_{600}$ readings were measured and means and SD's are presented (n = 3). (**E**) Strains lysogenic with phages SPBeta wt or SPBeta Δ*sar*$^{SPBeta}$ were complemented with either empty integration vector (pDG1664) or with construct containing *sar*$^{SPBeta}$ gene with its own promoter and SOS box. The number of phages produced in the presence or absence of MMC was quantified using 168 Δ6 as a recipient strain. The results are shown as PFUs ml$^{-1}$, with geometric means and SDs presented (n = 3). A two-tailed t-test on $\log_{10}$ transformed data was performed to compare mean differences. p values are indicated above the plots: $p < 0.01$ (∗∗).ns not significant

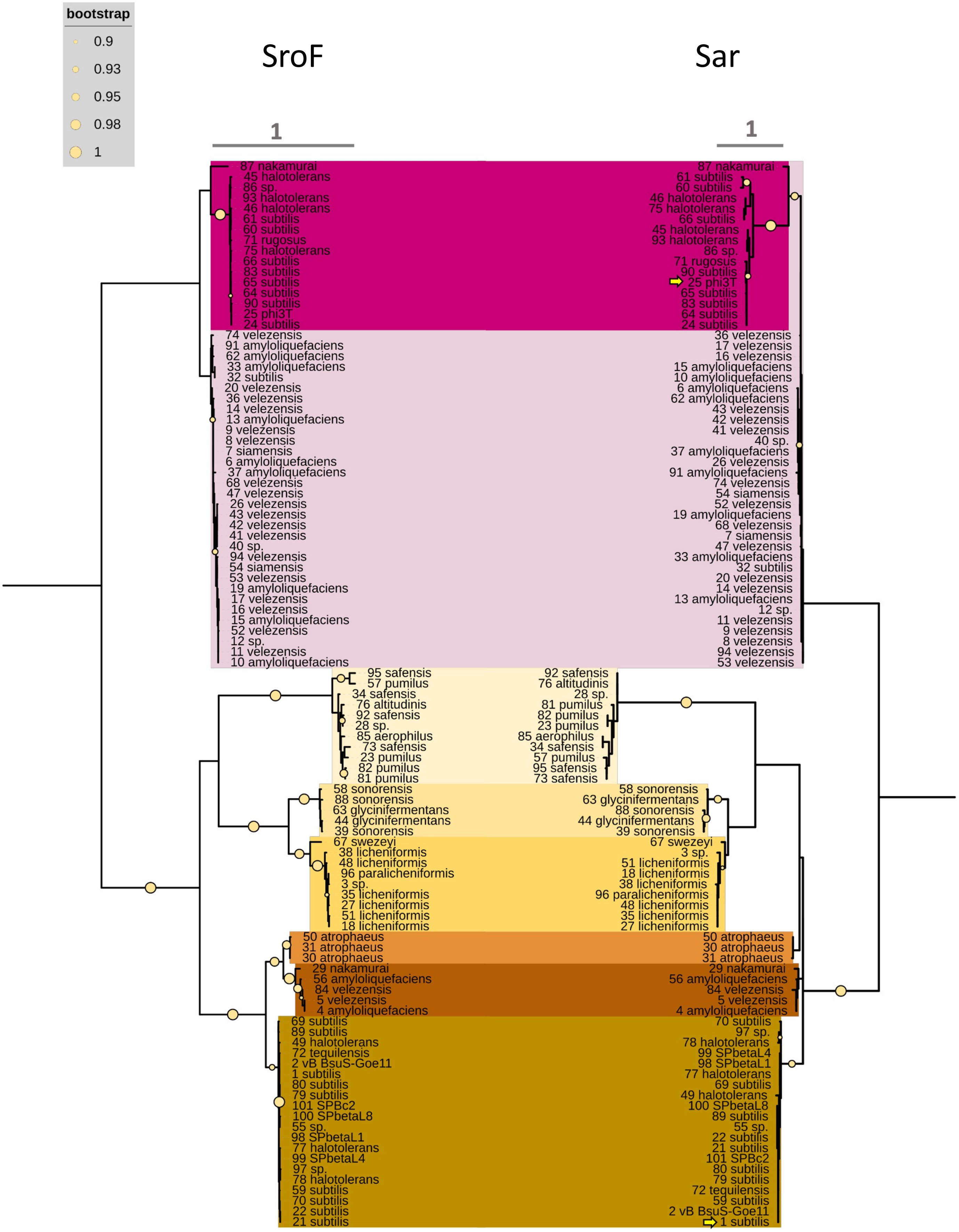


**Supplementary Figure S8: Phylogenetic trees based on the protein sequences of SroF and Sar, taken from 101 SPBeta-like phages, where at least one of these proteins is different than in other phages on the list**. Each leaf is labelled by index number in Supplementary Table S5 contains the species of the strain in which it was found (or phage name if it was sequenced separately). Colours mark major clusters appearing in both trees which are very similar. Circle sizes indicate bootstrap levels above 90% and lack of circle indicates a bootstrap confidence smaller than 90%. Phages Phi3T and SPBeta are highlighted by an arrow on both trees.

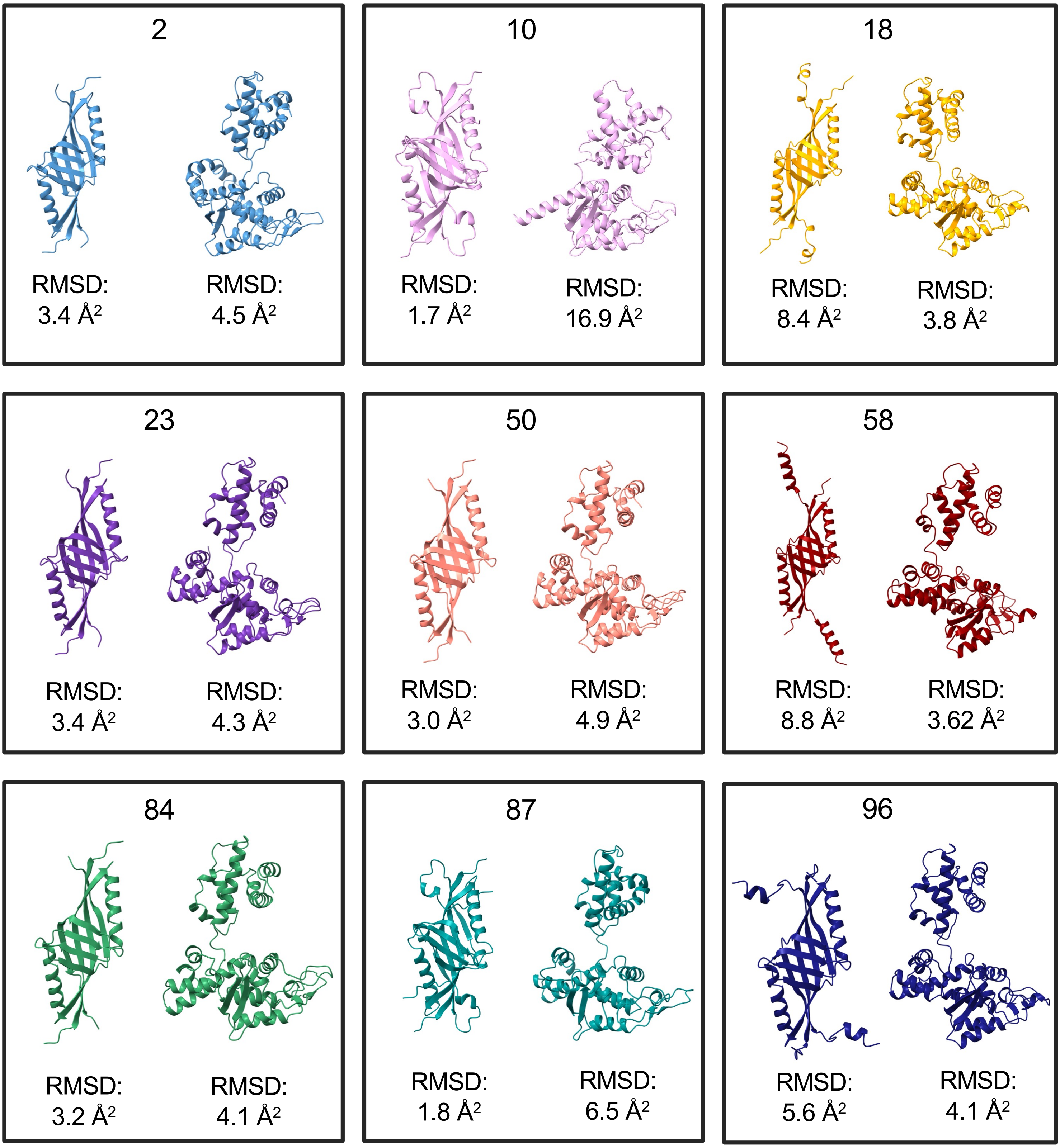


**Supplementary figure S9. Models for Sar and SroF found in SPBeta-like phages:** Ribbon representation of the obtained Alphafold models for the different homologs of both proteins. Number on top refers to the index number of the organism as presented in our supplementary table S5. RMSD values for the comparison of the full model with Sar$^{Phi3T}$ or SroF$^{Phi3T}$ (in the DNA bound conformation) are indicated below each predicted structure.
