## Supplementary Tables S1-S7 for "An unprecedented DNA recognition–mimicry switch governs induction in arbitrium phages": Supplementary Table S1.docx

**Supplementary Table S1. Data collection and refinement statistics**

|  | **SroF^Phi3T^-DNA** | **SroF^Phi3T^-Sar^Phi3T^** |
| --- | --- | --- |
| PDB code | 9RD5 | 9RGL |
| **Data Collection** |  |  |
| Wavelength (Å) | 0.9793 | 0.9793 |
| Space group | C 2 2 2_1_ | P 2_1_ 2_1_ 2_1_ |
| Cell dimensions |  |  |
| a, b, c (Å) | 75.2, 115.6, 154.4 | 78.9, 81.0,157.9 |
| α, β, γ (°) | 90, 90, 90 | 90, 90, 90 |
| Resolution | 48.82 - 2.5 (2.59 - 2.5) | 38.29 - 3.05 (3.16 - 3.05) |
| R*pim* | 0.018 (0.072) | 0.998 (0.912) |
| R*merge* | 0.063 (0.252) | 0.114 (1.164) |
| Mean I/ σ(I) | 28.43 (8.74) | 13.48 (2.68) |
| CC1/2 | 0.998 (0.992) | 0.998 (0.912) |
| Completeness (%) | 99.90 (99.79) | 99.46 (99.49) |
| Redundancy | 13.0 (13.3) | 9.6 (9.9) |
| **Refinement statistics** |  |  |
| Resolution | 2.5 | 3.05 |
| Number of reflections | 23650 (2327) | 19828 (1941) |
| *R*_work_ | 0.2069 (0.2978) | 0.2400 (0.3476) |
| *R_free_* | 0.2526 (0.3107) | 0.2819 (0.3924) |
| *CC*_work_ | 0.917 (0.850) | 0.957 (0.804) |
| *CC_free_* | 0.920 (0.821) | 0.909 (0.816) |
| N° non-hydrogen atoms | 3809 | 6615 |
| macromolecules | 3679 | 6594 |
| Ligand/ion | 9 | 0 |
| Solvent | 121 | 21 |
| B factors |  |  |
| macromolecules | 48.39 | 127.84 |
| Ligand/ion | 88.31 |  |
| Solvent | 49.97 | 66.54 |
| Rmsd bonds (Å) | 0.004 | 0.002 |
| Rmsd angles (°) | 0.73 | 0.46 |

Statistics for the highest-resolution shell are shown in parentheses.
