## Supplementary Tables S1-S7 for "An unprecedented DNA recognition–mimicry switch governs induction in arbitrium phages": Supplementary Table S2.docx

**Supplementary Table 2: SroF^Phi3T^ contacts with DNA and Sar^Phi3T^**.

| **SroF** | | **Contacts** | | |
| --- | --- | --- | --- | --- |
| Element | Residue | DNA nucleotide | DNA Backbone | Sar |
|  | Y5 |  |  | E33/K11*/M13* |
| α_1_ | R16 |  | T9 |  |
|  | Y17 |  | T9 |  |
|  | S19 |  | T10 |  |
| α_2_ | A21 | A11 | A11 |  |
| α_2_ | T22 | T10 | T9/T10 |  |
| α_2_ | L25 | T10/A11/A15* |  |  |
| α_2_ | Y26 | T9/T10 | T8/T9 |  |
| α_2_ | K29 | T16* |  |  |
| α_2_ | R31 |  | A14* |  |
| α_2_ | F35 |  |  | A44/V45 |
| α_2_ | R38 |  |  | E33 |
|  | D41 |  |  | M13* |
| α_4_ | D58 |  |  | K48 |
|  | K62 |  | A15*/T16* |  |
|  | S63 |  | T16*/A17* |  |
| α_5_ | E65 | T8/A17*/A18* | A17* |  |
| α_5_ | S66 | T16* | T16* |  |
| α_5_ | R68 | T7/T8 |  |  |
| α_5_ | R71 |  | T7/T8 |  |
| α_5_ | A72 | T8/T9 | T8 |  |
| α_5_ | Q76 |  | T8/T9 |  |
|  | Y95 |  | T7/T8 |  |
| α_6_ | D99 | T7 | T6/T7 |  |
|  | T112 |  |  | N52 |
|  | D143 |  | T14 |  |
|  | G144 |  | T13 |  |
| α_9_ | Y145 |  | T12/T13 | E82 |
| α_9_ | Q146 |  | T13 | D80 |
| α_9_ | H147 |  | T14 | D80 |
|  | K170 |  | T12 |  |
| β_3_ | Y195 |  | G7* | D96* |
| α_12_ | L197 |  | A6*/G7* | D96* |
| α_12_ | N198 |  | A6*/G7* | D96* |
|  | A202 |  | A6* |  |
|  | S203 |  | A6* |  |
|  | G204 |  | A5* |  |
|  | K205 | A24/T25/A2*/T4*/T5* | A24/T25/T3*/T4* |  |
|  | L206 | T22/A23 | A23/A24/A5* |  |
|  | R207 |  |  | E93* |
| β_4_ | F208 |  | A23/A24 |  |
| β_4_ | N210 |  | A23 |  |
|  | L220 |  | A8*/A9* |  |
|  | K221 |  | G7*/A8* | Y87*/E89* |
|  | Y222 |  | T22/A23 |  |
|  | K223 | T21/T22/G7* | A6*/G7*/A8*/T21/T22 |  |
|  | S224 |  | T21 |  |
|  | P225 |  | T21/T22 |  |
|  | D226 |  | T10* |  |
|  | Q227 |  | A8*/A9* |  |
|  | S228 |  | A9* |  |
|  | Q230 |  | A9* |  |
|  | D232 |  | A9* |  |
| α_13_ | K233 | T13/T14 |  | D21/S25* |
| α_13_ | F234 | T14/T15/T10*/A11* |  | Q19/G20/D21/S25* |
| α_13_ | L235 |  | A8* |  |
| α_13_ | H237 | T14/T15 | T13/T14/T15 | N76/D80 |
| α_13_ | R238 | A8*/A9* | G7*/A8*/A9* | Q19/Y87*/L99*/Y102* |
| α_13_ | S239 |  | A8* |  |
| α_13_ | K241 |  |  | D74/N76/E101* |
| α_13_ | T242 |  | G7* | S98*/L99* |
| α_13_ | Q244 |  | T15 | I75 |
| α_13_ | K245 |  |  | E97*/E100*/E101* |
| α_13_ | I246 |  |  | S98* |
|  | P250 |  | T15 |  |
|  | Y251 |  |  | N52/I75 |
|  | T253 |  | T14/T15 | E79 |
|  | P254 |  | T14 |  |
| α_14_ | K255 |  | T13/T14/A17*/A18* | E79 |
| α_14_ | N256 |  | A17* |  |
| α_14_ | N259 |  | A17* |  |
| α_16_ | V278 |  | G3/T4 |  |
| α_16_ | K282 |  | G3 |  |
|  | K296 | A2/G3/T4/A23* | A2 |  |
|  | F297 | G3/T4 | G3 |  |
|  | A298 | T4 | T4 |  |
|  | S299 |  | T4 |  |
| α_17_ | Q300 | A5/A6/T7/A20* | A5 |  |
| α_17_ | R303 |  | T4/A5 |  |
| α_17_ | K304 | T8/A18* |  |  |

Interactions through the same SroF residue for the DNA and Sar are highlighted with a yellow background. Asterisks indicate interaction with the second DNA strand or second monomer in Sar homodimer. Residues in red highlight salt bridge interactions.
