## Supplementary Tables S1-S7 for "An unprecedented DNA recognition–mimicry switch governs induction in arbitrium phages": Supplementary Table S3.docx

**Supplementary Table S3. Phage Phi3T proteins identified in SroF^Phi3T^ pull-down assays**

| **Sample ID** | **Gene** | Protein Description |
| --- | --- | --- |
| 1 | phi3T_134 | Uncharacterized protein |
| 2 | phi3T_3 | Thermonuclease like protein |
| 1 | phi3T_97 (SroF^Phi3T^) | Core-binding (CB) domain-containing protein |
| 2 | phi3T_97  (SroF^Phi3T^) | Core-binding (CB) domain-containing protein |
| 1 | phi3T_99 | Uncharacterized protein |
| 2 | phi3T_99 | Uncharacterized protein |
