## Supplementary Tables S1-S7 for "An unprecedented DNA recognition–mimicry switch governs induction in arbitrium phages": Supplementary Table S4.docx

| **Strain** | **Mutation** |
| --- | --- |
| JP26025 | Additional A in aimR binding site (underlined) (AAGTTCCAGAAATTCAAAAATCAAAAAA**A**TAAGAACAT) |
| JP26026 | Additional A in aimR binding site (underlined) (AAGTTCCAGAAATTCAAAAATCAAAAAA**A**TAAGAACAT) |
| JP26027 | 1 nt deletion in nt position 560/1137 of *aim*R gene, leading to frameshift. |
| JP26028 | 1 nt deletion in nt position 203/1137 and point mutation (A🡪G) in position 206/1337 of *aim*R gene, leading to premature stop codon. |

**Supplementary Table 3: Identified mutations from evolution experiments with phage Phi3T Δsar^Phi3T^ pDR110 sar^Phi3T^ pAX01 sar^Phi3T^ induced with IPTG and xylose. Sequencing results for survivors.**
